# High reelin expression can explain why the entorhinal cortex is a cradle for Alzheimer’s disease

**DOI:** 10.1101/2025.06.04.655278

**Authors:** Asgeir Kobro-Flatmoen, Jagir R. Hussan, Peter J. Hunter, Stig W. Omholt

## Abstract

The entorhinal cortex (EC) plays a crucial role in memory functions. Long before the clinical symptoms of Alzheimer’s disease (AD) emerge, it has already undergone significant degeneration, making it a primary site for the onset of the disease. The reasons for this remain elusive. It was recently shown that in layer II neurons of the anterolateral entorhinal cortex (alECLII neurons), which are especially prone to display a very early increase in intracellular amounts of amyloid-*β* peptide (A*β*) and hyperphosphorylated tau protein (p-tau), the large glycoprotein reelin binds to A*β*, suggesting that reelin functions as a sink for intracellular A*β*. The expression of reelin is extraordinarily high in alECLII neurons compared to most other cortical neurons. Here, we show by computational modeling that, in a senescent physiology predisposing to frequent inflammation-driven A*β*42 production bursts, the intracellular amount of A*β*42-reelin complexes can accumulate to extraordinarily high levels in alECLII neurons compared to the vast majority of cortical neurons. This explains experimental data showing that intracellular accumulations of A*β*42 positive material ranged from 20 to 80% of the total cytoplasmic volume in EC neurons from patients with sporadic AD. We also show that this extreme intracellular aggregation can cause the accumulation of detrimental hyperphosphorylated tau fragments. Thus, when exposed to recurrent AD-promoting stress, the exceptionally high expression of reelin in alECLII neurons appears to be instrumental in their early demise relative to other cortical neurons.

## Introduction

Long before the appearance of clinical symptoms in the course of Alzheimer’s disease (AD), increased intracellular amounts of A*β* followed by p-tau are predominantly restricted to neurons of layer II (LII) of the entorhinal cortex (EC), especially those of the anteriolateral domains of EC (alEC) (Kobro-Flatmoen et al., 2021; Berron et al., 2021). Multiple studies show that alECLII neurons are typically among the first to degenerate in the course of the disease (Gómez-Isla et al., 1996; Kordower et al., 2001; Kulason et al., 2019; Holbrook et al., 2020). And a recent study, where tau-positron emission tomography scans of 1,612 individuals were analyzed using an algorithm that combines disease progression modeling with traditional clustering to achieve probabilistic spatiotemporal partitioning and classification, identified four distinct but stable spatiotemporal subtypes of tau accumulation in AD consistent with the conception that all initially emerged from the EC (Vogel et al., 2021). Since alECLII neurons are integral to the generation of spatial and relational representations on which our memory system depends, it explains why the initial symptoms of most, if not all types of AD include impaired memory and links the most typical early degenerative events in AD to the deprivation of specific functionally characterized neurons.

More than 20 years ago, it was convincingly shown by D’Andrea et al. (D’Andrea et al., 2001, 2002) that (i) A*β*42 accumulated selectively as discrete granules associated with lysosomes and/or their derivatives in the soma of principal neurons in the EC of AD brains (72-78 years), (ii) the intracellular volume of A*β*42 positive material ranged from 20-80% of total cytoplasmic volume, (iii) there was a very strong linear relationship (*R*^2^ = 0.87) between the size of extracellular amyloid plaques and the size of adjacent neurons, and (iv) regions that were abundantly populated with principal neurons displaying excessive A*β*42 accumulations also contained evidence of neuronal lysis. To the best of our knowledge, these findings have not been experimentally refuted. Instead, a solid amount of data accumulated in the past 20 years shows that lysosomal dysfunction is a commonly observed abnormality in proteinopathic neurodegenerative diseases (Koh et al., 2019). And recent findings in mice (Lee et al., 2022) suggest that there is in fact a causal link between a dysfunctional endosome-autophagosome-lysosome pathway (EALP), pronounced intraneuronal accumulation of A*β*, and loss of neuronal structural integrity leading to cell death and subsequent plaque formation.

The diameter of the 4.5 kDA A*β*42 molecule is approximately 2 nm (Walsh et al., 1997; Erickson, 2009). If we, based on the available data (Kobro-Flatmoen and Omholt, 2025), take the soma volume of an alECLII neuron to be approximately 5000 *µm*^3^, and assume that A*β*42 peptides are packed rather loosely (about 1/3 of what can be obtained by random close packing of spheres), it follows that the copy number of A*β*42 must be about 1×10^11^ to occupy 50% of the soma volume. Suppose that this number has accumulated over 10 years and that there has been no decay, the mean production rate must have been approximately 1.1×10^6^ molecules per hour. Since this figure is unrealistically high (Hausser et al., 2019), it strongly suggests that to reach the observed soma volume occupancy of A*β*42 positive granules in EC cells (D’Andrea et al., 2001), A*β*42 must have accumulated together with one or more much larger molecules.

The function of the large glycoprotein reelin in adult mammalian brains includes synaptic modulation, induction of enhanced spine density, and promotion of long-term potentiation (Qiu et al., 2006). Most alECLII neurons are unique among cortical excitatory neurons by expressing extraordinarily high levels of reelin (dubbed Re^+^alECLII neurons in the following). It was recently shown that reelin and A*β* most likely make a stable complex (A*β*42_*reelin*_) (Kobro-Flatmoen et al., 2023), and since the volume of the 388 kDa reelin molecule is approximately 86 times greater than that of an A*β*42 molecule, this provides a possible mechanism by which excessive amounts of A*β*42 positive granules can accumulate in Re^+^alECLII neurons if they are exposed to frequent bursts of A*β*42 production induced, for example, by inflammation (Franceschi et al., 2018; Walker et al., 2019; Stranahan and Mattson, 2010).

Here, using a computational model based on and integrating a wide range of experimental data, we show that in the context of a senescent physiology predisposing to frequent inflammation events that trigger A*β*42 production (Welikovitch et al., 2018; Kinney et al., 2018), the intracellular amount of A*β*42_*reelin*_ in Re^+^alECLII neurons can easily accumulate to occupy 50% of the volume of the soma, while there is no marked accumulation of A*β*42_*reelin*_ in cortical neurons expressing low levels of reelin (LR neurons). Furthermore, through impairment of reelin signaling by A*β*, we demonstrate that it is fully conceivable that this accumulation is causally related to the observation that high intracellular levels of aggregation-prone p-tau fragments and cell death appear much earlier in Re^+^alECLII neurons than in other cortical neurons. Intriguingly, simply by reducing the affinity between reelin and A*β*42, the model recapitulates the phenotype of a rare mutation in the gene coding for reelin, recently shown to cause extreme resistance to AD (Lopera et al., 2023).

Together, the results indicate that a major reason why EC is a cradle for AD is the extremely high intra-cellular accumulation of A*β*42_*reelin*_ in Re^+^alECLII neurons. And they further support the opinion that such accumulation can lead to lysis of neurons (D’Andrea et al., 2001; Gouras et al., 2000; Lee et al., 2022).

## Computational model

### Outline of the model

The basic component of the computational model is a differential equation model of the relevant dynamics in a single cortical neuron. The model describes the per hour rate of change of the intracellular copy numbers of free A*β*42 ([A*β*]), free reelin ([Reelin]), A*β*42-reelin complexes tagged for degradation by the EALP ([A*β*_*reelin,L*_], A*β*42 similarly tagged for degradation by the EALP [A*β*_*L*_)]), phosphorylated GSK3*β* prevented from inducing tau hyperphosphorylation ([GSK3*β*_*p*_]), free hyperphosphorylated tau ([*tau*_*p*_]), and aggregation-prone p-tau fragments associated with the EALP ([*tau*_*p,agg*_]). The model is designed to capture the effect of increasing the production rate of A*β*42 from a normal baseline level to a markedly higher level for a given number of hours before allowing the production rate to return to the baseline level. This mimics the effect of the onset and end of a short-term inflammatory event (Walker et al., 2019).

To predict the long-term time evolution of the seven state variables over [Ω_1_, Ω_2_] years caused by recurrent EC inflammation events of varying durations and of multifactorial peripheral or central nervous system origins that characterize senescent physiology (Franceschi et al., 2018; Furman et al., 2019; Walker et al., 2019; Zhang et al., 2023; Mekhora et al., 2024), we run the dynamic model according to the information provided by an inflammation event vector *I*. This vector was created by a stochastic process that, based on published statistics on the prevalence of inflammatory events, defined when an inflammation event occurred and how long it lasted. For simplicity, we let the probability of an inflammation event in a given month in [Ω_1_, Ω_2_] increase linearly from 0 to Ψ, and let its duration be sampled uniformly from the set {*D*_*min*_, ‥, *D*_*max*_} days (see Fig. S1 in the Supplementary Information for further details). During an inflammation period, the production rate of [A*β*] was increased to a maximum constant value (*α*_*infection*_), otherwise it was kept at a constant baseline value (*α*_*baseline*_). Unfortunately, quantitative data on the variability of the duration of brain inflammation events as a function of age that could guide the parameterization underlying *I* do not appear to be available. However, based on data from the literature and statistics available on the Web site of the US Government Centers for Disease Control and Prevention, we can conservatively estimate that, on average, a person will be exposed to around 70-75 brain inflammation events from 50 to 72 years (see the supplementary text “Data motivating the parameterization of the inflammation vector *I*” accompanying Fig. S1 for details). This estimate was used to parameterize Ψ. All other parameter values of the dynamic model were kept constant throughout the 22-year period.

The computational model was coded in Python in a Jupyter lab environment. The SciPy Python library (Virtanen et al., 2020) was used to generate the time evolution of the differential equation system in the period [Ω_1_, Ω_2_] (scipy.integrate.solve_ivp with the Radau solver). The code producing the results and the figures in the main text and the Supplementary Information is available on Zenodo (link to be provided).

### The dynamic model

The dynamic model is an extended version of a previously published model (Kobro-Flatmoen and Omholt, 2025) that focused on the short-term dynamics of [A*β*], [Reelin],[A*β*_*reelin*_], and [GSK3*β*_*p*_]. Since available data make it plausible that A*β* and p-tau are integral parts of an evolutionary very old intraneuronal immune response to acute microbial or viral infection of the mammalian brain (Soscia et al., 2010; White et al., 2014; Bourgade et al., 2015; Soscia et al., 2010; Vandamme et al., 2012; Kumar et al., 2016a; Fulop et al., 2018; Butler and Walker, 2021; Wainberg et al., 2021; Li et al., 2021), the previous model was designed to capture key elements of this response in native physiology. The dysfunctional operation of this response in a senescent inflammation-prone physiology may be etiologically important in the early development of AD in EC (Kobro-Flatmoen and Omholt, 2025). However, the results of the model presented here are not dependent on this being the case. Specifically, the cause(s) of the inflammation event(s) is not decisive, as the model only assumes that an inflammation event is capable of triggering an increase in A*β*42 production and that the maximum production rate is substantially higher in Re^+^alECLII neurons than in LR neurons (Kobro-Flatmoen and Omholt, 2025).

The dynamic model is given by Eqs. [1]-[7] below. To facilitate understanding, the functions [8]-[11] contained in four of the seven equations are listed separately.

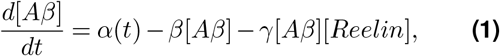

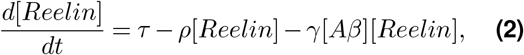

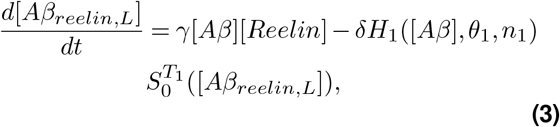

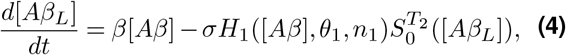

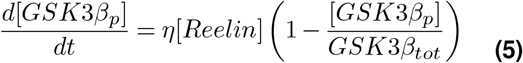

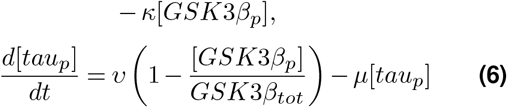

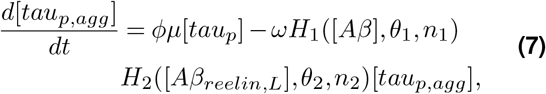

where

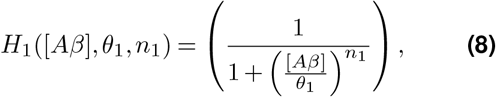

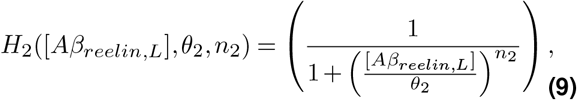

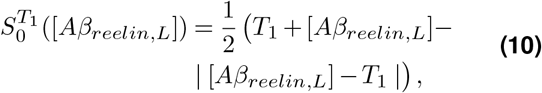

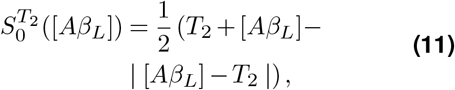

The premises underlying the above model are:

***Equation [1]***. The production rate of A*β*42 (*α*(*t*)) is at a very low constitutive baseline level (*α*_*baseline*_) or at a much higher constitutive level when the cell is exposed to an inflammation event (*α*_*infection*_). The degradation of A*β*42 is a first-order process (*β*[A*β*]), and there is a second-order binding reaction between A*β*42 and reelin (*γ*[A*β*][Reelin]). We assume a 1:1 binding of the reelin to A*β*42.

***Equation [2]***. Reelin production is constitutive (*τ*), a first-order process (*ρ*[*Reelin*]) labels free reelin for degradation, and the second-order binding reaction between A*β*42 and the reelin is identical to the last term in Eq. [1]. Reelin labeled for degradation might, under certain conditions, accumulate intracellularly (see the descriptions of Eqs. [3] and [4] below). However, since this is unlikely to be functionally significant in an AD context, we have not considered this possibility.

***Equation [3]***. The subscript L in [*Aβ*_*reelin,L*_] indicates that the A*β*42_*reelin*_ complex is rapidly tagged for degradation by the EALP. In native physiology, it is reasonable to assume that the degradation can be approximated by first order kinetics (Kobro-Flatmoen and Omholt, 2025). However, in senescent physiology, this is unlikely to be the case, since the disruption of the EALP consistently appears as an early and progressive characteristic of the pathophysiology of AD (Knopman et al., 2021), leading to accumulation of degradation-tagged cellular cargo. A solid body of data implicates that dysfunction of the lysosomal vacuolar (H+)–adenosine triphosphatase (v-ATPase) proton pump is of particular significance (Song et al., 2020). Both A*β* and *β*-C-terminal fragment (*β*-CTF), another cleavage product of the amyloid precursor protein (APP), appear to be capable of altering v-ATPase to cause lysosomal alkalinization leading to a general decrease in lysosomal degradation (Koh et al., 2019; Lee et al., 2022; Im et al., 2023). Since the expression level *β*-CTF is likely to be highly correlated with the expression level of APP, which is closely related to the expression level of its proteolytic metabolite A*β* (Rovelet-Lecrux et al., 2006; Ozmen et al., 2008; Wu et al., 2019), we used only [A*β*] in the inverse Hill function [8] to express that when the levels of A*β* and *β*-CTF pass a critical threshold, the processing capacity of the EALP starts to decline rapidly (see Fig. S2).

The function [10] expresses the assumption that the degradation of A*β*42_*reelin,L*_ is a first order process *δ*[*Aβ*_*reelin,L*_] up to a given copy number threshold *T*_1_. When the number of A*β*_*reelin,L*_ complexes reaches *T*_1_, the processing machinery becomes saturated, so the number degraded per hour becomes constant (*δT*_1_) (Fig. S3). This is likely to be a conservative assumption since it is conceivable that as the number of EALP bodies associated with accumulation increases, this may lead to a demand for v-ATPase that is beyond what its production machinery has evolved to handle, causing alkalinization and thus an even stronger decline in degradation capacity. Note that the equation assumes that the saturation kinetics also applies when there is no A*β*/*β*-CTF driven reduction in the processing capacity of the EALP. Smooth versions of the functions [10] and [11] (Mahdi Ghazaei Ardakani and Magnusson, 2018) were used in the numerical simulations as required by the numerical solver used (see the caption of Fig. S3 for further details).

***Equation [4]***. The processing rate of intracellular A*β*42 tagged for degradation in the EALP ([A*β*_*L*_]) (Albaret et al., 2023; Wiersma et al., 2019; Kumar et al., 2016b; Zaarur et al., 2014) follows the same kinetics as that of A*β*42_*reelin*_ (see Fig. S3).

***Equation [5]***. By binding to its main receptor in the brain, the apolipoprotein E receptor 2 (ApoER2), reelin triggers a signaling cascade that strongly inhibits otherwise constitutively active glycogen synthase kinase 3*β* (GSK3*β*) (Hiesberger et al., 1999; Beffert et al., 2002; Ohkubo et al., 2003), which is one of the main kinases that phosphorylates tau. A major reason for our focus on this kinase is that a physical association between active GSK3*β* and tangle-like inclusions is initially observed in ECLII neurons (Pei et al., 1999) and that interaction with A*β* reduces the capacity of reelin to inhibit GSK3*β* kinase activity (Cuchillo-Ibáñez et al., 2013). The equation expresses the assumption that the phosphorylation rate of the serine 9 (S9) epitope on GSK3*β*, whose phosphorylation causes its inhibition (Tanji et al., 2002), is a function of the reelin level and the total copy number of S9-unphosphorylated GSK3*β* (GSK3*β*_*tot*_) (*η* is a constant) and that the dephosphorylation rate is a first order process (*κ*[GSK3*β*_*p*_]). See reference (Kobro-Flatmoen and Omholt, 2025) for further justification of the phosphorylation term.

***Equation [6]***. The rate of p-tau production is proportional to the amount of S9-unphosphorylated GSK3*β* and *υ* is the maximum production rate. The rate at which p-tau is marked for degradation in the EALP (Lee et al., 2013) is a first order process (*µ*[*tau*_*p*_]). In a normal mature neuron, the tau concentration is estimated to be approximately 2 *µ*M and practically all tau is assumed to be bound to microtubules (Iqbal et al., 2010). The protein is recovered in three main states (soluble, oligomeric, and fibrillized) from the brains of AD subjects, and although there is as much normal cytosolic tau in the AD brain as in the normal aged brain, the level of abnormally hyperphosphorylated tau is several times higher and as much as 40% of the tau is oligomeric (Iqbal et al., 2010). This suggests that the intracellular level of normal tau is under homeostatic feedback control and that p-tau is not part of this regulation, which supports the use of the simple production term in Eq. [6].

***Equation [7]***. The EALP appears to contribute both to the production of proaggregating forms by the fragmentation of tau and to the clearance of tau aggregates (Wang et al., 2009). The reduced translocation of the F1 tau fragment across the lysosomal membrane appears to be a critical mechanism to promote the formation of tau oligomers on the surface of these organelles through the creation of the aggregation-prone fragments F2 and F3 by cathepsin L lysosomal protease (Wang et al., 2009; Boyarko and Hook, 2021). The production term in Eq. [7] indicates that a fraction *ϕ* ≤ 1 of *tau*_*p*_ tagged for EALP degradation is on the surface of lysosomes converted to truncated forms of p-tau that are prone to aggregation [*tau*_*p,agg*_]. Since tau fragments are generated via chaperone-mediated autophagy, they can be produced even if lysosomal fusion is blocked by lysosomal alkalinization (Murrell-Lagnado and Frick, 2019) inasmuch as lysosomal proteases remain operative (Funk and Kuret, 2012). This suggests that the production term in Eq. [7] also applies during A*β*/*β*-CTF driven lysosomal alkalinization.

The function *H*_2_(.) ([9]) in the degradation term expresses the assumption that there is an inverse sigmoidal relationship between the amount of lysosomal A*β*42_*reelin*_ and the rate at which the truncated forms of p-tau are degraded. Hence, the lysosomal uptake rate of the truncated p-tau forms will in the non-inflammatory case be according to first order kinetics (*ω*[*tau*_*p,agg*_]) until the number of EALP bodies containing large amounts of A*β*42_*reelin*_ starts to predominate (Fig. S4).

### Parameterization of the differential equations

The parameter values used are listed in Table 1. The parameter values that differ in two types of neurons are highlighted by light blue shading. See reference (Kobro-Flatmoen and Omholt, 2025) for a justification of the values used for *α*_*baseline*_, *α*_*infection*_, *β, γ, τ, ρ, δ, η, GSK*3*β*_*tot*_, *κ* and soma volume.

**Table 1.**
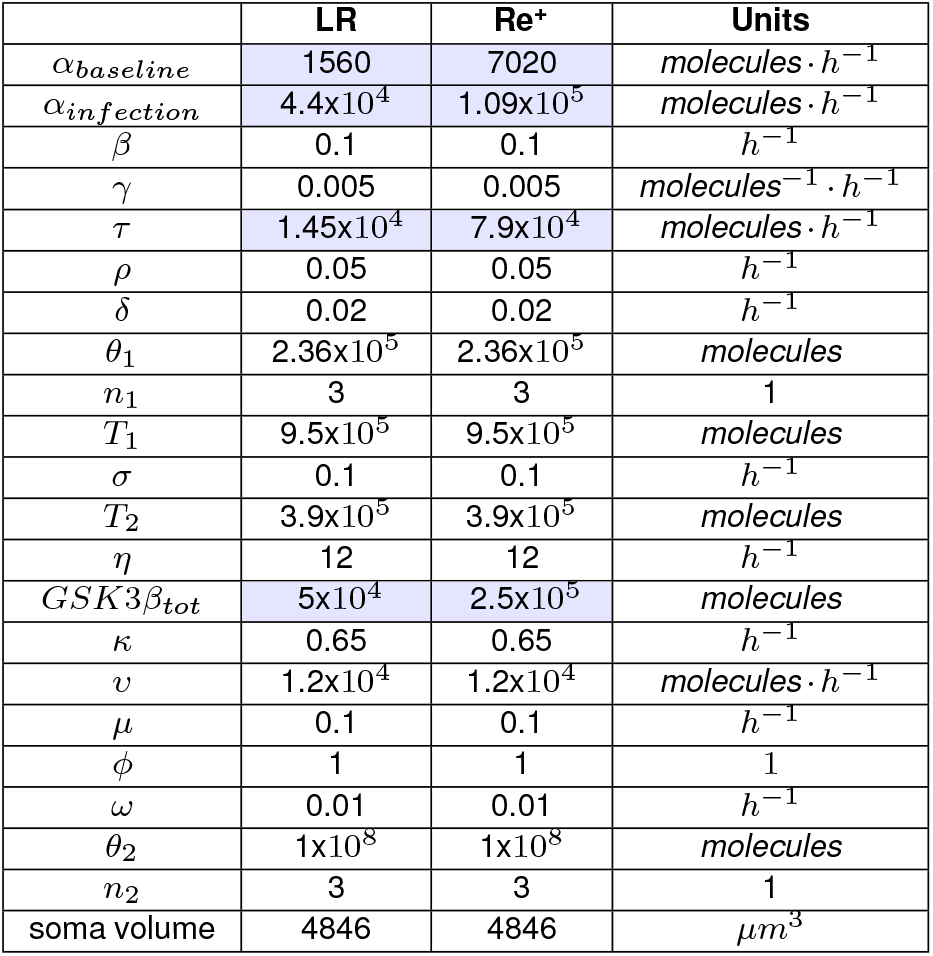
Parameter values used to describe LR and Re^+^alECLII neurons in the dynamic model.

We let *θ*_1_ be approximately 80% of the maximum cellular concentration of free A*β*42 during an inflammation event. The value of *n*_1_ reflects that the effect of the A*β*/*β*-CTF driven alkalinization of lysosomes is assumed to be a moderate sigmoidal process. The thresholds *T*_1_ and *T*_2_ in functions [10] and [11] were allowed to be approximately 50% above the steady state copy number of [A*β*_*reelin,L*_] and [A*β*_*L*_] in LR neurons during an inflammation event in native physiology, assuming no effect of lysosomal alkalinization. Since there is little evidence for accumulation of A*β*42 and p-tau in EALP under normal conditions in native physiology, we let *σ* = *µ* = *β*. The value of *υ* reflects the assumption that the neuron can increase its production of p-tau quite quickly when reelin is not inhibiting the GSK3*β* pathway. Due to reduced translocation of the F1 tau fragment across the lysosomal membrane, we let *ω* be 1/10 the value of *µ*. The value of *n*_2_ assumes the existence of a moderate sigmoidal dose-response relationship. The value of *θ*_2_ assumes that the copy number of A*β*_*reelin,L*_ has to be about ten times higher than the number experienced by an LR neuron in native physiology exposed to an inflammation event lasting a few days before it starts to influence the degradation rate of *tau*_*p,agg*_.

## Results

### Dynamic response to a short-term inflammation event

The predicted dynamics of six of the seven state variables associated with a single short-term inflammatory event in a LR and a Re^+^alECLII neuron are shown in Figs. S5 and S6. The main distinctions between the two cases are that in the latter, the levels of [A*β*_*reelin*_] and [A*β*_*reelin,L*_] become approximately six times higher and that [A*β*_*reelin,L*_] decreases much slower after the cessation of the inflammation event. The parameter values that are different in the two cases are the A*β* production rates *α*_*baseline*_ and *α*_*infection*_, the constitutive production rate of reelin (*τ*), and the copy number of GSK3*β*_*tot*_. These differences are justified in reference (Kobro-Flatmoen and Omholt, 2025).

### Long-term dynamic response to recurrent inflammation

By running the full computational model where the A*β*42 production (Eq. [1]) was successively evoked for a given time period according to the vector of inflammation events (*I*) as defined above, we obtained the development of the seven state variables over a 22-year span (Ω_1_ = 50,Ω_2_ = 72) in an LR and a Re^+^alECLII neuron (Figs. 1 and 2, only four variables are shown).

**Figure 1.**
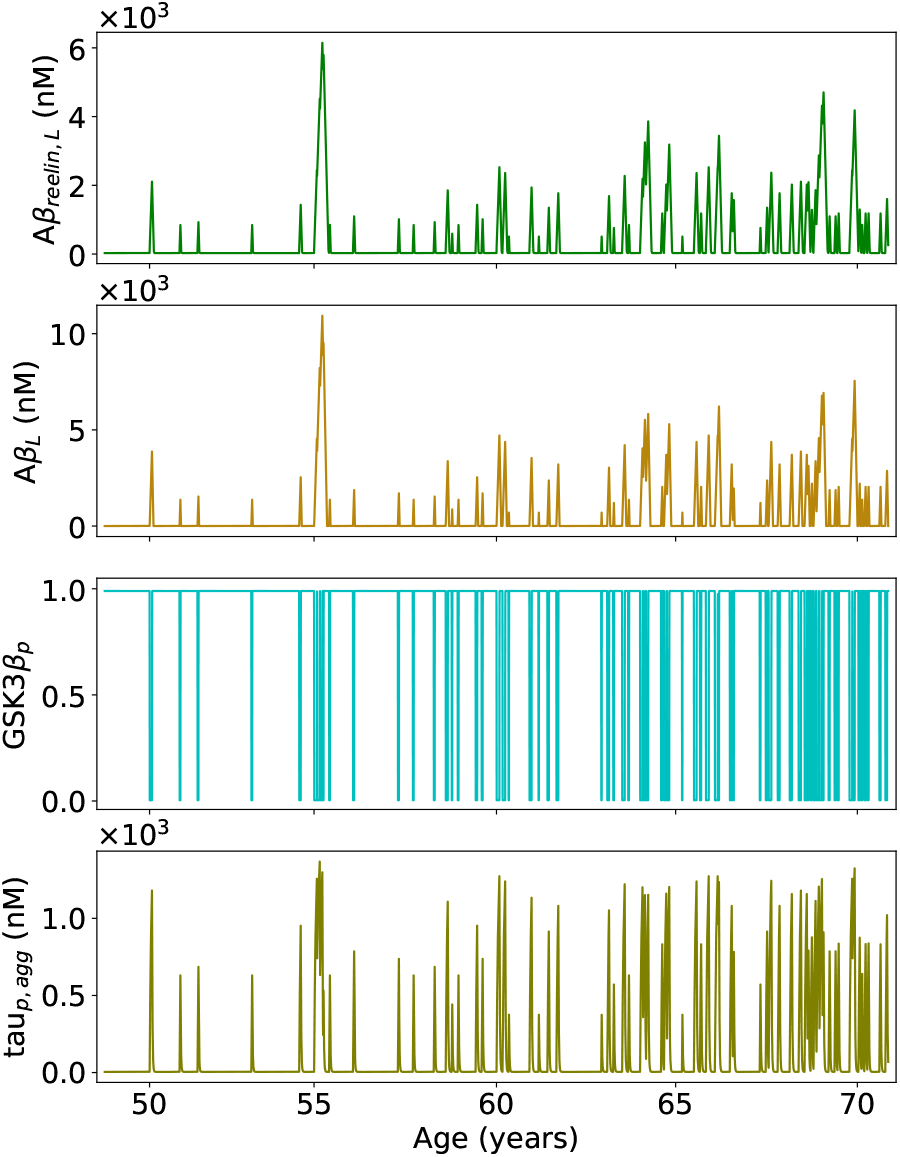
Predicted time evolution of four of the seven state variables in an LR neuron over 22 years. GSK3*β*_*p*_ is expressed as proportion of the total number of GSK3*β*. The parameter values used to generate the inflammation event vector *I*: Ω_1_ = 50 years, Ω_2_ = 72 years, Ψ = 0.6, *D*_*min*_ = 5 days, *D*_*max*_ = 29 days.

**Figure 2.**
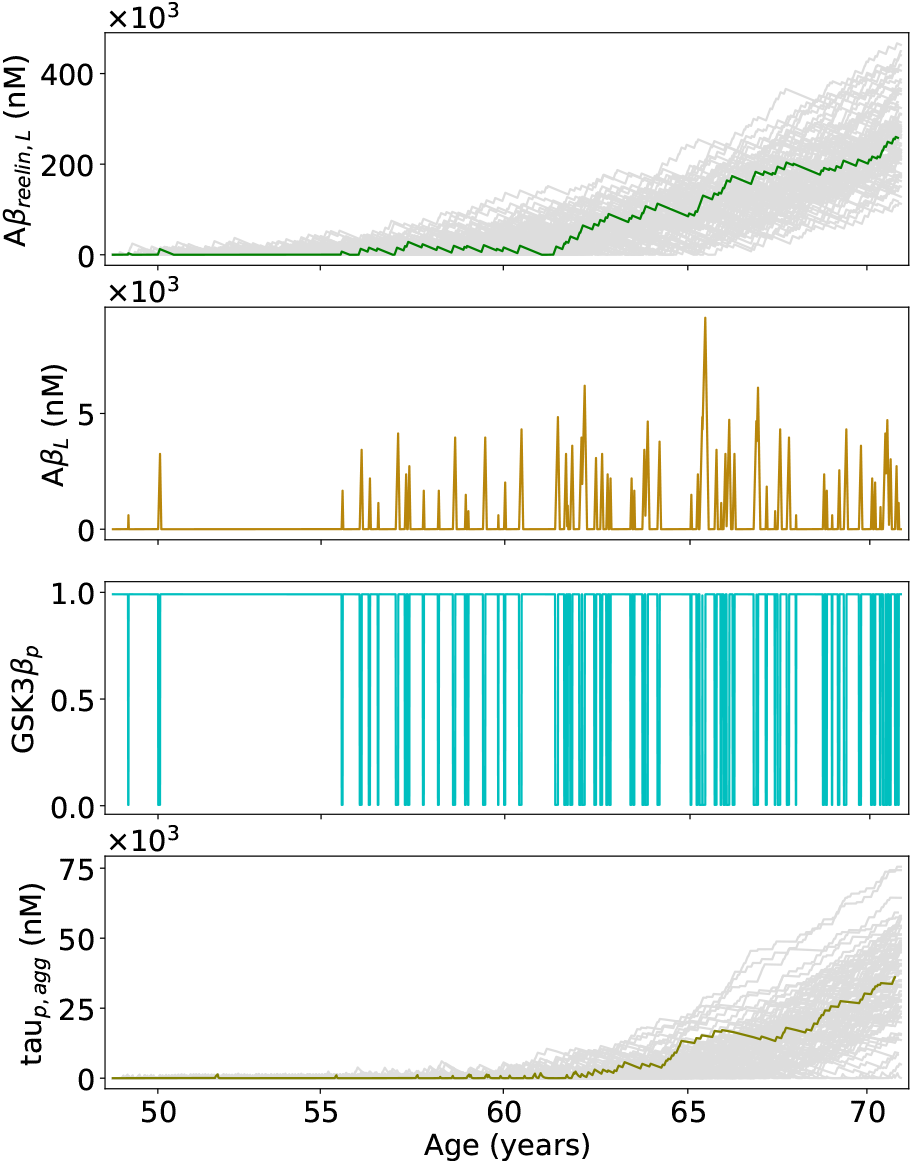
Predicted time evolution of four of the seven state variables in a Re^+^alECLII neuron over 22 years. *I* is identical to the one used for the LR neuron (Fig. 1).

In an LR neuron (Fig. 1), the model predicts that the intracellular concentration [A*β*_*reelin,L*_], [A*β*_*L*_] and [*tau*_*p,agg*_] will increase during each inflammation event, but that degradation between each bout will be large enough to prevent the steady accumulation of the three species. The proportion of GSK3*β* being phosphorylated at S9 [GSK3*β*_*p*_] is predicted to be ≈ 1 between inflammation events and ≈ 0 during an inflammation event.

In the Re^+^alECLII neuron case (Fig. 2), the model predicts that the time series pattern of [A*β*_*L*_] and [GSK3*β*_*p*_] will be similar to that of an LR neuron, while [A*β*_*reelin,L*_] and [*tau*_*p,agg*_] will start to accumulate steadily at a given time point. Since *I* is a stochastic vector, we have illustrated the spread in the accumulation patterns of [A*β*_*reelin,L*_] and [*tau*_*p,agg*_] obtained from using 100 additional versions of *I* based on the same parameterization as the green and olive lines in the first and fourth panels (light gray lines). We see that the time development of the two state variables can vary substantially depending on the inflammation history given by *I*, but the model consistently predicts that [A*β*_*reelin,L*_] and [*tau*_*p,agg*_] will accumulate intraneuronally in Re^+^alECLII neurons but not in LR neurons.

It should be noted that [A*β*_*reelin,L*_] and [*tau*_*p,agg*_] can also accumulate in an LR neuron if we dramatically increase the number and mean duration of inflammation events. However, the model then predicts that the soma volume occupancy of [A*β*_*reelin,L*_] in a Re^+^alECLII neuron will be much higher than 100%.

The volume of an A*β*42_*reelin*_ complex is at least 470 *nm*^3^ assuming a density of 1.37 g/*cm*^3^ (Erickson, 2009). We do not know how tightly A*β*42_*reelin*_ complexes can pack in EALP bodies. Assuming that they are packed about a third as densely as can be obtained by a random close packing of spheres, it follows that a soma volume occupancy of 50% would require the concentration of A*β*42_*reelin*_ to be about 375 *µM*. Thus, the model appears fully capable of recapitulating the experimental data reported by D’Andrea et al. (D’Andrea et al., 2001, 2002) (Fig. 2, first panel).

The extreme accumulation of A*β*42 positive material in EC neurons of AD subjects (D’Andrea et al., 2001) implies that the mean degradation rate (and export) of this material must have been substantially lower than the mean production rate for a long period of time. If we exchange the saturation function 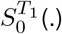 with [A*β*_*reelin,L*_] and keep all remaining parameters fixed, we do not get any accumulation of A*β*_*reelin,L*_ in the Re^+^alECLII neurons. This supports the notion that degradation must begin to follow zero order kinetics when the A*β*_*reelin,L*_ level reaches a given threshold, preventing the return to the baseline level in non-inflammatory periods and leading to accumulation of A*β*_*reelin,L*_.

Although the results must be interpreted with caution due to the paucity of experimental data that could have further constrained the choice of parameter values, they clearly indicate that enhanced constitutive reelin expression in Re^+^alECLII neurons makes these neurons much more vulnerable to the onset of AD than LR neurons. As shown in the following, this contention is supported by recent experimental data.

### The effect of reduced affinity between A*β*42 and reelin

A presumably rare mutation in the reelin gene (*RELN*) has recently been shown to cause extreme resistance to EC degeneration in an individual with the autosomal dominant familial AD (ADFAD) PSEN1-E280A mutation (Lopera et al., 2023). The individual remained cognitively intact until age 67, while carriers of PSEN1-E280A usually show clinical AD symptoms in their mid to late 40s. The reelin mutation clearly prevented EC neurons from developing a severe tau tangle burden, despite being exposed to a high A*β* load (Lopera et al., 2023).

One possible explanation is that the mutation causes a dramatic reduction in the affinity between A*β*42 and reelin. Letting the value of the parameter *γ* (Eqs. [1], [2], and [3]) become much less in a Re^+^alECLII neuron than that used to produce Fig. 2 (assuming the reelin mutation is dominant), while keeping all other parameters fixed and not including the effect of the PSEN1-E280A mutation, the model predicts that there will be marginal accumulation of A*β*42_*reelin*_ and aggregation-prone p-tau fragments (Fig. 3, panels 1 and 3). In contrast, it predicts that a large amount of [A*β*_*L*_] can accumulate (Fig. 3, panel 2). [GSK3*β*_*p*_] is predicted to be *≈* 1, regardless of whether there is an infection event or not (not shown). Since the model is clearly capable of recapitulating key phenotypic effects of this reelin mutation, it appears worthwhile to determine the ability of the associated reelin-variant to bind to A*β*.

**Figure 3.**
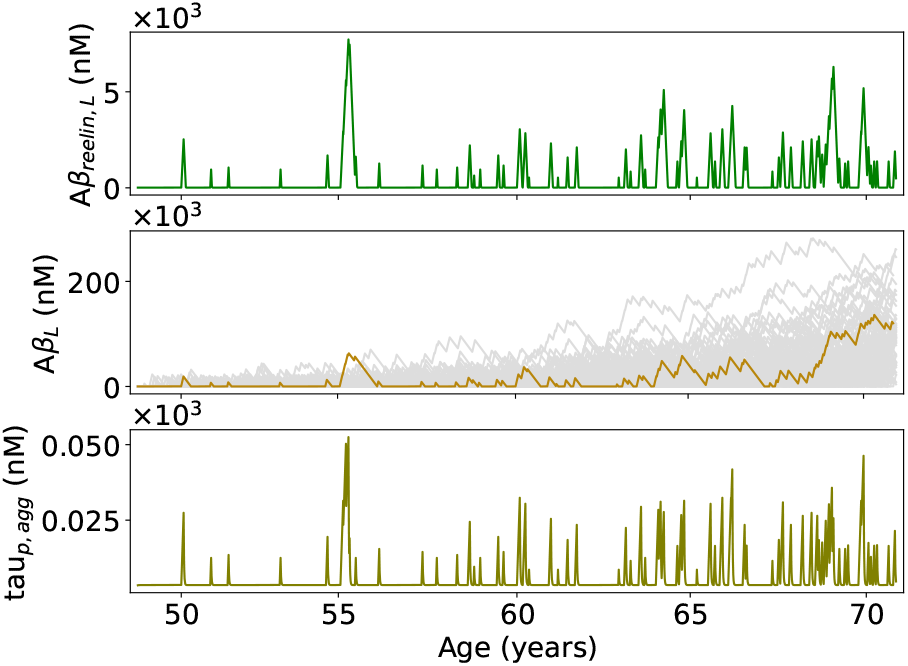
Predicted time evolution of three of the seven state variables over 22 years in a Re^+^alECLII neuron carrying a mutated *RELN* allele. Here, we let *γ* = 1*×* 10^−8^, while keeping all other parameters fixed. *I* is identical to the one used for the wild type Re^+^alECLII neuron (Fig. 2).

### Accumulation rate of A*β*_*reelin,L*_ in layer II vs. layer III neurons

Neuronal death in subjects developing AD probably starts several years earlier in ECLII than in ECLIII (Gómez-Isla et al., 1996; Kordower et al., 2001; Arendt et al., 2015). If this neuronal death is causally related to the soma volume occupancy of A*β*_*reelin,L*_, the accumulation of A*β*_*reelin,L*_ in ECLIII will have to be much slower than in ECLII. To test whether the model predicts this, we quantified the lateromedial expression levels of reelin in Re^+^ECLII and Re^+^ECLIII neurons in three 3 month and three 12 month old wild-type rats (Figs. S11 - S13; see captions for details of the experimental protocol and how the data were compiled to give relative reelin expression levels for all six animals combined). Assuming a linear relationship between the expression level of reelin and its production rate (*τ*), and taking advantage of the observation that the relative expression levels of reelin are approximately the same in humans (Pérez-García et al., 2001), we then used the model to predict the intragroup and intergroup variation of the accumulation rate of A*β*_*reelin,L*_ as a function of time.

The model predicts temporal accumulation patterns of A*β*_*reelin,L*_ that are consistent with the empirically observed temporal variation in neuronal cell death in EC (Kulason et al., 2018) (Fig.4). The soma volume occupancy is predicted to reach, for example, 25% in Re^+^ECLII neurons located toward the lateral extreme (i.e., close to the rhinal fissure) several years earlier than those located toward the medial aspect. And the first incidence of 25% soma volume occupancy in Re^+^ECLIII neurons is predicted to occur several years later than in Re^+^ECLII neurons. We acknowledge that this is an idealized calculation that does not account for variation in reelin expression within each of the 15 bins. If we include this variation, we will get a more heterogeneous situation, but the main prediction, concordant with solid empirical data showing that cell death will generally start in ECLII several years before it starts in ECLIII, will not be affected.

### Robustness of the *Aβ*_*reelin,L*_ accumulation results

Evidently, the degree of *Aβ*_*reelin,L*_ in Re^+^alECLII neurons depends on the parameters that define the generation of *I*. We quantified this dependency by sampling Ψ, *D*_*min*_ and *D*_*max*_ from respectively the sets {0.3, 0.4, 0.5, 0.6}, {1, 2, 3, 4} and {5, 10, 15, 20, 25, 30} in a full factorial design and recording the median intracellular *Aβ*_*reelin,L*_ concentration of 101 replicates at ≈ 72 years. An occupancy of the soma volume of 20% (≈ 150 *µM*) at 72 years is obtained when Ψ = 0.5 and *D*_*max*_ = 25 when *D*_*min*_ = 4 (Fig. S7). It should be noted that in the model we have not explicitly included any positive feedback effect on *Aβ*42 production rate from increasing levels of *Aβ*_*reelin,L*_ and *tau*_*p,agg*_. Hence, it is plausible that the number and duration of extra-cellularly driven inflammation events needed for severe aggregation of *Aβ*_*reelin,L*_ are less substantial than the above estimates suggest. When performing the same analysis on LR neurons, there is no sign of aggregation of *Aβ*_*reelin,L*_ (Fig. S8).

To study how sensitive the accumulation results in the two types of neurons were to variation of the parameters in the dynamic model, we generated 4608 sets of parameters following the Sobol sequence method (Saltelli et al., 2010) giving a discrepancy of 0.0007. The parameters *α*_*infection*_, *β, τ, ρ, δ, θ*_1_, and *T*_1_ varied between *±* 50% of the nominal values used to generate Figs.1 and 2. We let *γ* vary between [0.0005 − 0.05] and *n*_1_ was fixed to 3. We then generated 100 different inflammation profiles and Eqs. [1]-[3] were run for each set of parameters and each inflammation profile, resulting in 460,800 independent simulations. For each set of parameters, the maximum concentration of *Aβ*_*reelin,L*_ was determined for each inflammation profile. The median value among these 100 values was chosen as the representative *Aβ*_*reelin,L*_ value for that set of parameters. The simulation steps were repeated for the two sets of nominal parameters and the maximum, median and minimum *Aβ*_*reelin,L*_ values of the 100 simulations were determined. In Re^+^alECII neurons, the results show that for nearly all parameter sets the *Aβ*_*reelin,L*_ soma concentration reaches very high levels (Fig. 5, upper panel). In contrast, in LR neurons, almost all parameter sets give rise to a very modest accumulation of *Aβ*_*reelin,L*_. This indicates that the predicted marked difference in endpoint *Aβ*_*reelin,L*_ accumulation between the two neuron types is quite insensitive to parameter variation.

**Figure 4.**
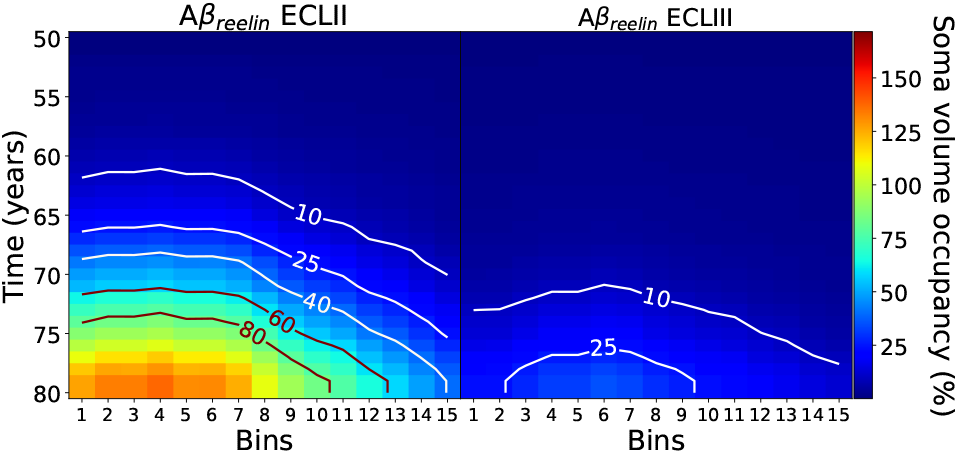
Predicted lateromedial soma volume occupancy of A*β*_*reelin,L*_ in ECLII and ECLIII as a function of time. Bin 1 corresponds to the lateral extreme position and increasing numbers refer to increasing medial positions. The *τ* value for Re^+^ neurons used to produce Fig.2 (Table 1) was multiplied with the estimated relative lateromedial reelin expression levels (divided into 15 bins) given in Fig. S13. All parameters were identical to those used to construct Fig. 2, except for the variation of *τ* imposed by the experimental data (Figs. S11-S13) and letting Ψ = 0.8 at 80 years. The soma volume occupancy is predicted to pass 100% in ECLII at about 75 years.

**Figure 5.**
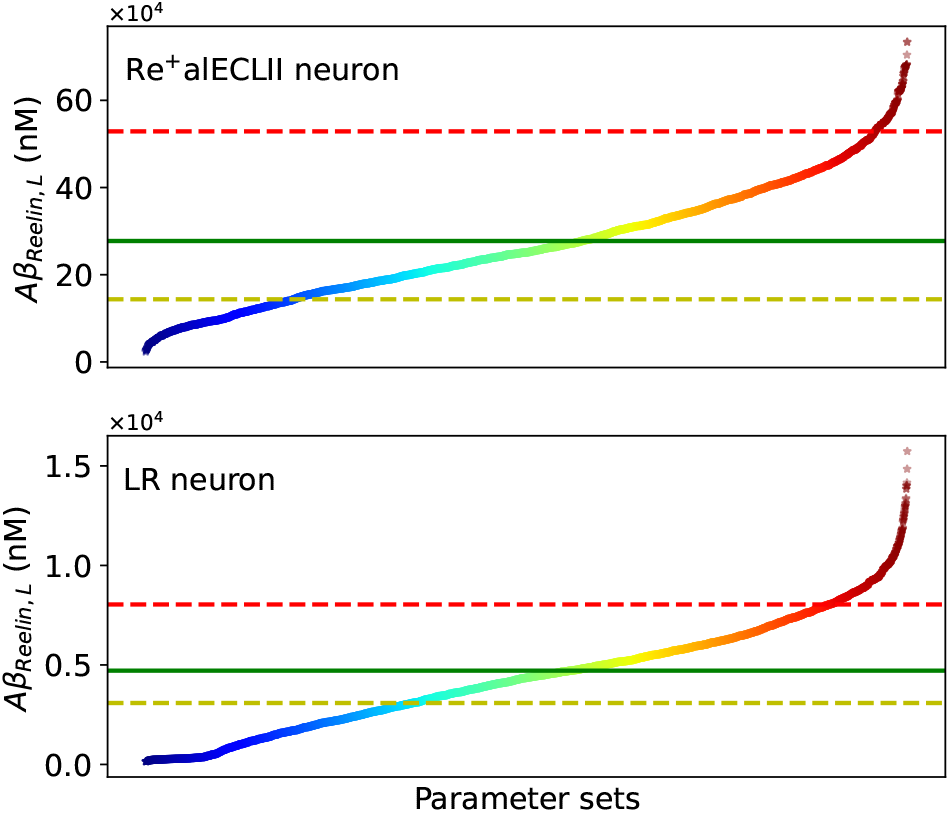
Median endpoint *Aβ*_*reelin,L*_ log soma concentration for multiparameter variations centered around the nominal parameter sets for the Re^+^alECII and LR neurons. The parameter sets are sorted in ascending order of their representative *Aβ*_*reelin,L*_ value. Horizontal lines show the maximum values (red dashed lines), median values (green solid lines), and minimum values (yellow dashed lines) values of *Aβ*_*reelin,L*_ for the nominal parameter sets. The corresponding soma volume occupancy percentages for the LR and Re^+^alECII neurons were (0.312, 0.502, 0.759) and (19.273,35.474,70.199), respectively. See main text for further details. We conducted two additional sensitivity analyzes using the open source Python library SALib v 1.47 (Herman and Usher, 2017), one estimating how much of the variance of the endpoint soma concentration of *Aβ*_*reelin,L*_ could be explained by the variation in each parameter (Fig. S9) and one that assessed the impact of the variation of parameters in Eqs. [5]-[7] on the endpoint [*tau*_*p,agg*_] value in Re^+^alECII neurons (Fig. S10). See captions of Figs. S9 and S10 for interpretations of the results.

## Discussion

Although we have succeeded in making quantitative predictions that are clearly in consonance with experimental observations, these predictions depend on several premises, some of which are not yet supported by conclusive experimental data. Of these, the most critical one is evidently the presumption that the A*β*42_*reelin*_ complex was the primary component in the soma-filling A*β*42 positive aggregates observed by D’Andrea et al. (D’Andrea et al., 2001). However, as already discussed, it is improbable that these lysosomal aggregates were made mainly of A*β*42. Since reelin is the only very large protein known to bind to A*β*42 that has markedly higher expression levels in alECLII neurons than in most other cortical neurons, it is clearly qualified as a candidate to make sense of the experimental data at hand (D’Andrea et al., 2001). Further support comes from the observation that A*β*42 accumulated in an insoluble pool within APP-expressing NT2N neurons differentiated from NT2 cells using retinoic acid (RA), while RAnaïve NT2 cells did not show any accumulation, despite expressing nearly equivalent levels of APP (Skovronsky et al., 1998). Since RA-induced differentiation of NT2 cells is accompanied by increased reelin expression (Chen et al., 2002), this suggests that the observed insoluble pool in NT2N neurons consisted of A*β*42_*reelin*_. In other cell culture studies reporting the accumulation of an insoluble A*β*42 pool associated with the EALP (Knauer et al., 1992; Yang et al., 1999) and where reelin apparently is not expressed at substantial levels, it is likely that the pool consists mainly of A*β*42.

Another critical presumption is the accumulation of aggregation-prone p-tau fragments in Re^+^alECLII neurons (Fig. 2), expressed by the function [11] in Eq. [7]. Currently, there is no direct experimental evidence for the hypothesized relationship between A*β*42_*reelin*_ and these tau fragments. However, due to the much earlier appearance of neurofibrillary tangles (NFTs) in Re^+^alECLII neurons than in most other cortical neurons (Kobro-Flatmoen et al., 2021; Berron et al., 2021), it follows that proaggregating p-tau fragments will precede the appearance of NFTs. Hence, it is not implausible that dysfunctional A*β*42_*reelin*_ filled lysosomes cause an increase in extralysosomal residence time of p-tau fragments when they exceed a certain number relative to functional lysosomes.

In addition to the above two concerns, several of the parameter values are ballpark estimates because of the paucity of experimental data. However, it should be noted that only Eqs. [1]-3] are needed to describe the evolution of [A*β*_*reelin,L*_], and all their parameter values, except those in functions [8] and [10], were used to describe the dynamics of A*β*42, reelin and A*β*42_*reelin*_ in native physiology (Kobro-Flatmoen and Omholt, 2025). Thus, they were deliberately not tuned to accommodate the quantitative correspondence between the predicted soma volume occupancy of [A*β*_*reelin,L*_] and the experimental data (Fig. 2).

We have deliberately addressed only the dynamics associated with A*β*42. This is of course a crude simplification as other A*β* variants are likely to be part of [A*β*_*reelin,L*_] and [A*β*_*L*_]. However, given the particular importance of A*β*42 in the development of AD, we consider this to be justifiable, especially as long as data that would allow increasing the resolution of the model are lacking.

D’Andrea et al. (D’Andrea et al., 2001) provided ample evidence for neuronal lysis as a source of plaques in EC. More recent findings in mice (Lee et al., 2022) indicating that cellular lysis can be a generic mechanism of plaque formation support this, although the matter is still debated. Although we do not know what the main triggers for neuronal lysis are, there is a well-documented strong link between the presence of p-tau oligomers and mitochondrial dysfunction (Shafiei et al., 2017; Szabo et al., 2020). And Re^+^ECLII neurons are clearly more prone to experience mitochondrial impairment very early in disease development than LR neurons (Terada et al., 2020; Olajide et al., 2021). One possible explanation for this is that since Re^+^ECLII neurons appear to have a very high energy demand (Omholt et al., 2024), they could be particularly vulnerable to a drop in energy supply. Our model predictions are clearly consistent with this scenario and the observation that NFT typically first appear in Re^+^alECLII neurons. However, our results are also consistent with the notion that a very high occupancy of the soma volume by EALP cargo can by itself trigger a cascade of pathophysiological processes, including mitochondrial dysfunction and formation of p-tau oligomers, which eventually lead to cell death.

If we assume that neuronal lysis is causing plaque generation in EC, we are confronted with the puzzle that although Re^+^ECLII neurons accumulate a conspicuously high amount of intraneuronal A*β*, ECLII forms far fewer plaques compared to the deeper layers of EC as AD progresses. Considering that Re^+^alECII neurons start to disappear well before mild cognitive symptoms begin to appear, a possible explanation is that dead cells, including those that have been lysed and may have formed transient plaques, are removed relatively quickly in the very early phase of AD development. This conception is supported by the observation that dead neurons are removed quite quickly by concerted operation of astrocytes and microglia (Damisah et al., 2020), and that microglia are capable of completely removing amyloid aggregates over a few weeks by digestive exophagy (Jacquet et al., 2024). Since our results indicate that the accumulation of A*β*42_*reelin*_ reaches critical levels in Re^+^ECLII neurons several years before Re^+^ECLIII neurons (Fig. 4), this suggests that the fully operational astrocyte/microglia system that is likely to exist in the very early phase of disease development has the capacity to remove cells that have been lysed without leaving behind a large number of plaques. However, as the disease progresses, this system will probably be prone to saturation effects and other types of dysfunctionalization.

Although our model ought to be looked upon as a first step towards a genuine representation of the dynamics causing the EC to be a cradle for AD, its capacity to make sound quantitative predictions indicates that such a representation is within reach if its construction is given sufficient theoretical-experimental attention.

## Acknowledgments

We thank Goran Simic for valuable comments on a previous version of this paper. This work was supported by the K. G. Jebsen Foundation, the Kavli Institute for Systems Neuroscience Centre of Excellence Grant, the Liaison Committee between the Central Norway Regional Health Authority (RHA) and the Norwegian University of Science and Technology (NTNU), and the Neurology Department, St. Olavs Hospital, Trondheim, Norway. The authors wish to acknowledge the use of New Zealand eScience Infrastructure (NeSI) high-performance computing facilities, consulting support and/or training services as part of this research. New Zealand’s national facilities are provided by NeSI and funded jointly by NeSI’s collaborator institutions and through the Ministry of Business, Innovation & Employment’s Research Infrastructure Programme. URL https://www.nesi.org.nz.

## Supplementary Information

**Figure S1.**
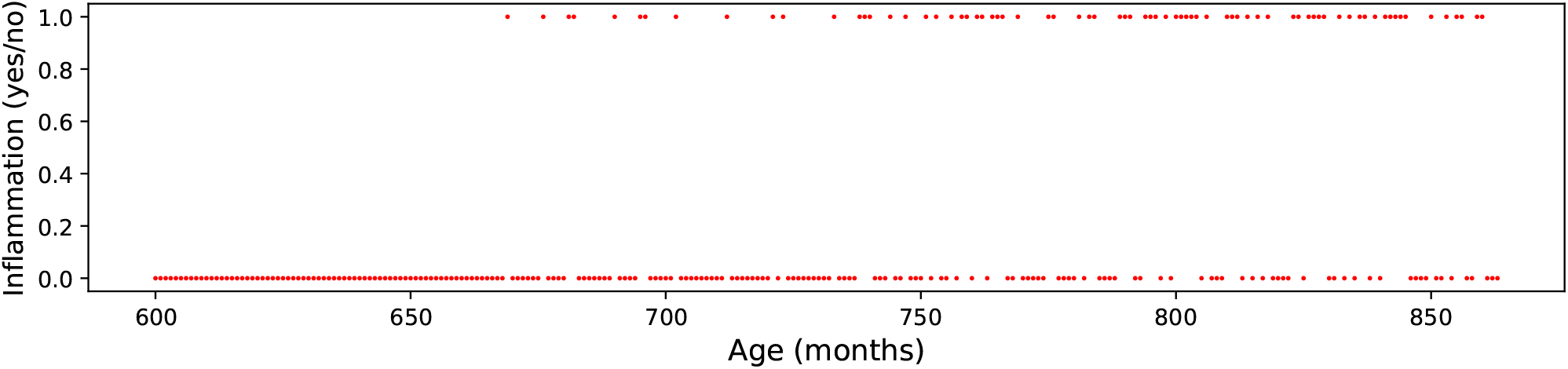
Distribution of inflammation events over the period [Ω_1_, Ω_2_] years. The distribution was generated by dividing the period into the corresponding number of months (*N*_*m*_). We then defined an inflammation propensity function *P* (*t*), *t* ∈{1, 2, …, *N*_*m*_ *}*. For simplicity, we let *P* (*t*) increase linearly, with *P* (1) = 0 and *P* (*N*_*m*_) = Ψ. We then sampled, for each month, a random number from *X* ∼ *U* (0, 1). If the number was ≤ *P* (*t*), then the neuron experienced an inflammation event starting on the first day of the month, otherwise not. The duration of an inflammation period was allowed to be *X* ∼ *U* (*D*_*min*_, *D*_*max*_), where *D*_*min*_ and *D*_*max*_ denote the minimum and maximum number of days that the inflammation event could last. The above procedure generated a vector (*I*) consisting of time-labeled elements describing when and for how long a neuron was exposed to an inflammation event during the period [Ω_1_, Ω_2_]. The parameter values used to generate the inflammation event vector *I* partly depicted here: Ω_1_= 50 years, Ω_2_ = 72 years, Ψ=0.6, *D*_*min*_ = 5 days, *D*_*max*_ = 29 days. This particular realization of *I* had 74 inflammation events with a mean duration of 16.8 days.

### Data motivating the parameterization of the inflammation vector *I(t)*

It is well known that several common infections and certain invasive medical procedures that trigger systemic inflammation also cause secondary neuroinflammation, and that bouts of neuroinflammation tend to become more detrimental to brain health as we age (Heneka et al., 2025). To arrive at a reasonable and conservative estimate of the number of times a person between the ages of 50-72 years experiences a neuroinflammatory episode as a result of systemic inflammation, we used statistics from the US Centers for Disease Control and Prevention (CDC) along with data from several published papers.

In the following, we provide estimates for the incidence rates of 12 of the most common causes of systemic inflammation in the relevant age range of 22 years. Causes are listed in order of estimated prevalence. We recognize that several of the causes of systemic inflammation listed below will influence each other and that this likely gives rise to complex interactions that cannot be captured by a single number. However, for the purpose of reaching a conservative estimate, we think that our approach is justifiable.

### Exposure to pollution and toxins

Environmental pollution is a major contributor to neuroinflammation, and is likely to disproportionately affect elderly people with a compromised blood-brain barrier (Block and Calderón-Garcidueñas, 2009; Costa et al., 2019). People in urban areas will experience several bouts of systemic inflammation per year, mainly due to air pollution (Meng et al., 2021; Thompson et al., 2010). Given that 80% of all people in the western world live in cities (UN_urban), we can safely assume that people will, on average, experience more than one bout of systemic inflammation due to exposure to pollution and toxins each year. We therefore, conservatively, set this number at 1.1 bouts / year, giving an estimate of 24 bouts of systemic inflammation due to exposure to pollution and toxins within our age range.

### Sleep deprivation

There are convincing data to show that sleep deprivation induces neuroinflammation (Cao et al., 2024). More than one-third of people aged 50 to 72 years are sleep deprived according to the CDC (that is, less than 7 hours of sleep per day) (CDC_sleep). Although person-to-person variation is likely to be high, we believe it is safe to assume that, on average, a person between the ages of 50-72 experiences at least one bout of substantial sleep deprivation per year. Thus, we reach 22 neuroinflammatory bouts caused by sleep deprivation in this age range.

### Vaccines

Although vaccines rarely cause serious complications and have saved approximately 154 million lives in the last 50 years alone (Shattock et al., 2024), they, as part of their effect, cause an inflammatory response that is likely to include a neuroinflammatory component (Nath, 2023). The recommended schedule for vaccination for people over 50 years old in the US, which is similar to that in most other countries in the western world, includes an annual influenza vaccine, at least two COVID-19 vaccines, plus other vaccinations against common pathogens such as, for example, diphtheria, tetanus, and whooping cough (CDC_vaccines). However, since not all people follow the recommended vaccination schedule, we conservatively estimate the number of neuroinflammation bouts due to vaccinations received between the ages of 50-72 to be 10.

### Allergic rhinitis (a.k.a. hay fever)

Allergic rhinitis involves a systemic inflammatory reaction and, although many details have yet to be worked out, such reactions also include a substantial neuroinflammatory component (Konstantinou et al., 2022). Data from the US show that about 7.8% of people have allergic rhinitis and that these people will experience between 3-10 inflammatory bouts each year; taking a conservative approach, we assumed that these people will experience 4 bouts. Thus, taking the number of people in the U.S. around time these statistics were collected (330 million) we may calculate that, on average, a person will experience 6.86 bouts of neuroinflammation due to allergic rhinitis over a 22-year period. Several other less common allergic reactions are also likely to result in a systemic (and hence neuro) inflammatory reaction. However, since we do not have data on their combined prevalence, and in the vein of aiming for a conservative estimate, we have not included these here.

### Major surgery

Major surgery induces systemic inflammation that is generally proportional to the invasiveness of the procedure (Watt et al., 2015; Lindholm et al., 2015; Bain et al., 2023; Lassig et al., 2019). Data from CDC, available for the year 2010 (CDC_surgery; see also CDC_procedures), show that the number of major surgeries in the U.S. population older than 45 years was 18.67 million. Since the number of people over 45 years of age in the US is 139 million, the average person is estimated to experience 2.86 bouts of neuroinflammation due to surgery over our target 22-year period.

### Influenza

The CDC estimates that, in the US, the number of influenza cases per year ranges from 9.3 to 41 million (CDC_-fluburden). Using the mean (25 million), and with a US population of about 330 million at the time when these statistics were collected, the average person is estimated to experience 1.67 bouts of neuroinflammation due to influenza over a 22-year period.

### Norovirus and other gastrointestinal diseases

Based on CDC data (CDC_norovirus), about 20 million US citizens get infected by norovirus per year, while about 4.4 million citizens will suffer at least one bout of some other gastrointestinal illnesses per year. With a population of about 330 million at the time when these statistics were collected, the average person may be expected to experience 1.6 bouts of neuroinflammation due to such illnesses over a 22-year period.

### Respiratory Syncytial Virus (RSV)

RSV tends to cause serious inflammation in small children and older adults, and has an annual incidence rate for the elderly of about 7% (Kim and Choi, 2024). Between ages 50-72 we can thus calculate that, on average, an individual will experience (22 × 0.07) 1,54 bouts of RSV.

### Parainfluenza Virus

Parainfluenza virus is known, among other conditions, to cause bronchitis and pneumonia in older adults (Pawełczyk and Kowalski, 2017). CDC data show that the annual U.S. incidence rate is approximately 5% (CDC_incidence), which for a population of 330 million gives an estimate of 1.1 neuroinflammatory bouts over the 22-year period.

### Urinary tract infection

Excluding the childhood period, biomarker data from the US and Canada indicate that women will experience at least 1-2 bouts of urinary tract infections over a lifetime, while for men the number is less than 1. However, the number of self-reported incidents is much higher (Foxman, 2010). Based on these data, we assumed one bout of neuroinflammation due to urinary tract infection during the 22-year period.

### Human Metapneumovirus (hMPV)

Like parainfluenza virus, hMPV is a common virus that tends to cause serious inflammation in small children and older adults (Shafagati and Williams, 2018). CDC data show that the annual incidence rate is approximately 3.8% (CDC_hMPV). This gives an estimate of 0.84 neuroinflammatory bouts over the 22-year period. Other common upper respiratory tract infections involve, for example, adenovirus (annual incidence rate of about 3%) and rhinovirus (annual incidence rate of approximately 20%) (CDC_rhino). Since these are believed to only rarely cause a substantial inflammatory response, we have chosen to exclude them from the estimate.

### Inflammatory Bowel Disease (IBD)

The US incidence rate of IBD in subjects older than 50 years is approximately 1% (Lewis et al., 2023), giving an average estimate of 0.2 bouts of neuroinflammation due to IBD over the 22-year period.

### Conclusion

Our estimates apply to people in the Western world, although they are likely to be also relevant to people in many nonwestern countries. Based on data on the 12 sources of systemic inflammation listed above, we find that, conservatively estimated, a person between ages 50-72 will experience on average >70 bouts of systemic inflammation that is likely to lead to secondary neuroinflammation in individuals in the given age range. Due to the lack of quantitative data, we have not included neuroinflammation events caused by poor oral health and changes in the gut microbiome (Heneka et al., 2025). Furthermore, in senescent physiology, recurrent inflammation events of peripheral and central nervous system origin and of variable duration may be caused by several other processes in addition to those listed here (Franceschi et al., 2018; Furman et al., 2019; Walker et al., 2019; Zhang et al., 2023; Mekhora et al., 2024). In line with the above considerations, the inflammation vector *I(t)* depicted in Fig. S1 contains 74 inflammation events of varying duration. Due to stochasticity, this number will vary somewhat for each realization of *I(t)*, but will overall be in the range of 65-80 events. Unfortunately, we have not found any data that can be used to constrain the parameter values that define the mean length and the variability of the duration of an inflammation event. In Fig. S1, the mean duration is 16.8 days. This may be an overestimate, but considering that the total number of inflammation events may very well be higher than what we have stipulated, allowing the mean duration to be reduced, we think that the parameterization of *I(t)* used generates a family of inflammation profiles that are physiologically sound.

**Figure S2.**
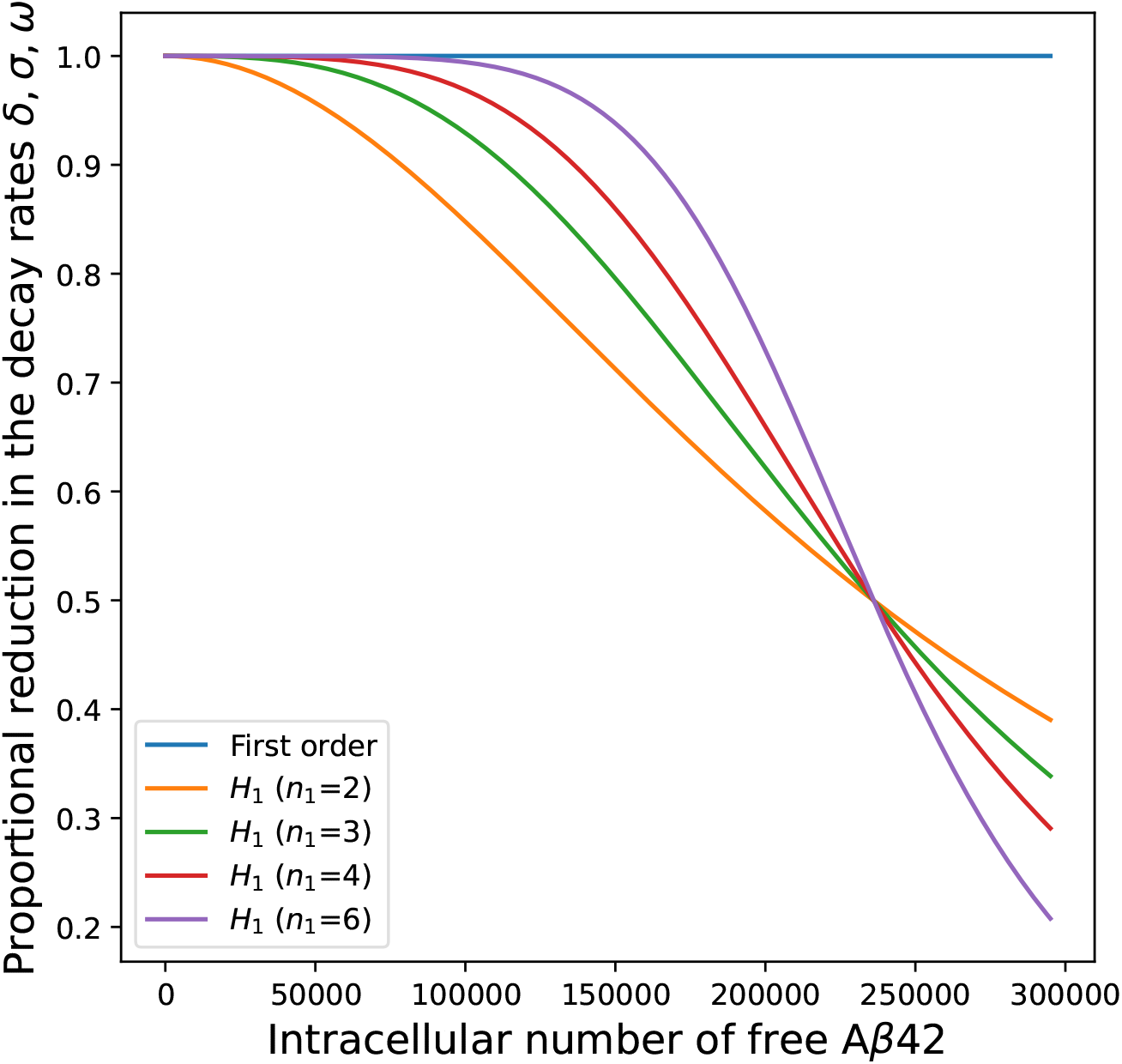
Examples of function [8] (*H*_1_([*Aβ*], *θ*_1_, *n*_1_)) that is part of Eqs. [3], [4] and [7]. The blue line refers to a first order process, i.e. the degradation rate is given by the specific decay constant times [*Aβ*]. Since we assume the existence of a moderately sigmoidal dose-response relationship, we chose to use the green inverse Hill function graph with *n*_1_ = 3. *θ*_1_ = 2.36×10^5^ molecules, i.e. 80% of the maximal copy number of free A*β*42 during an inflammation event when *H*_1_(.) is set to zero.

**Figure S3.**
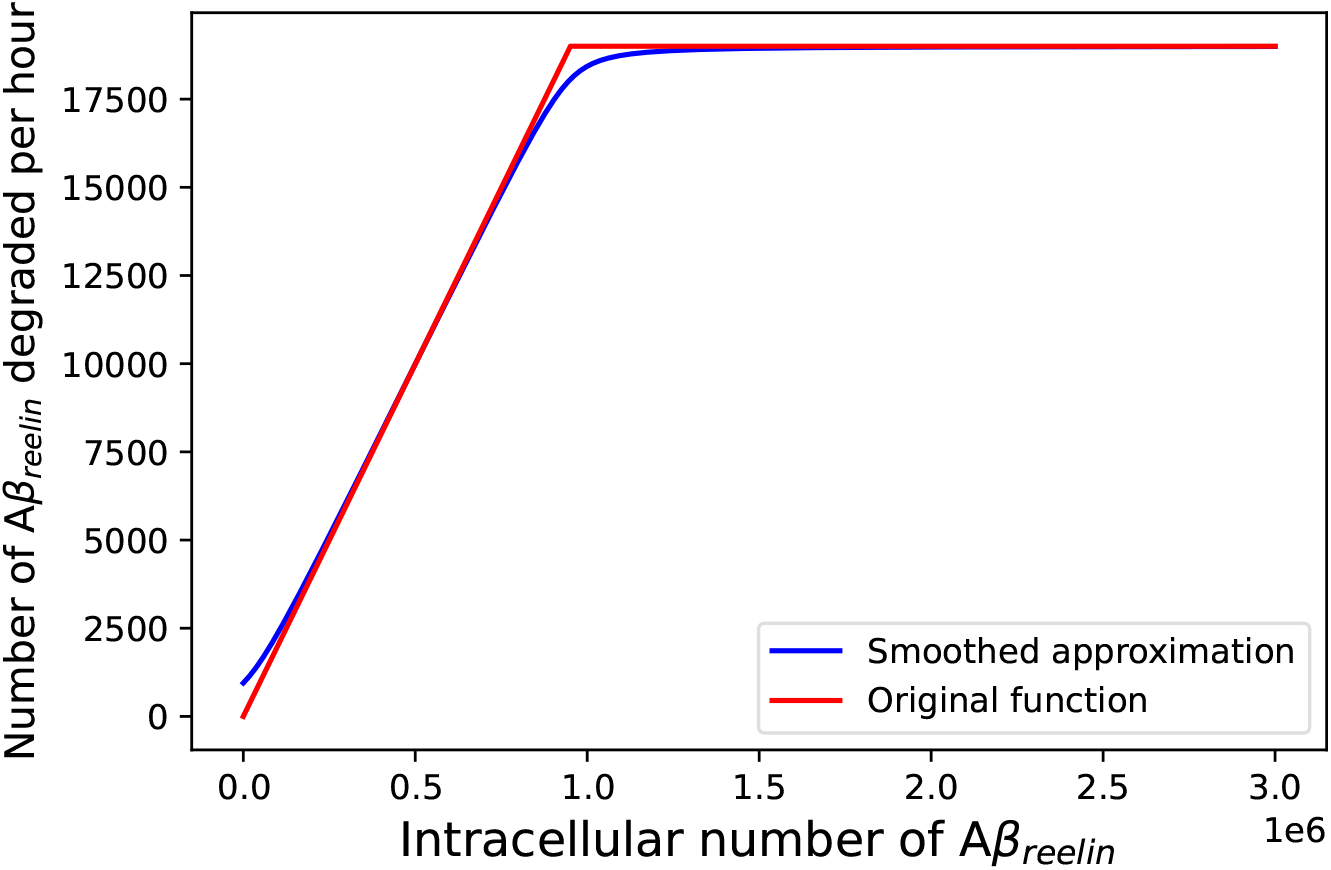
Smooth version of the threshold function [10] 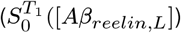 used in Eq. [3]. The function (red graph) expresses that as long as the intracellular copy number of *Aβ*_*reelin,L*_ is below *T*_1_, the degradation rate is according to a first order process (*δ* [*Aβ*_*reelin,L*_]). When it is above, the degradation kinetics becomes zero order due to saturation of the degradation machinery. Since this function is not smooth as required by the numerical solver, in the simulations we instead used the smooth version: 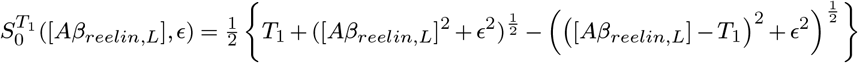. The blue graph depicts 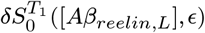 with *ϵ* = 1*x*10^5^ in order to make the graphs easily distinguishable. In the simulations we let *ϵ* = 1, making the graphs indistinguishable. The values of *δ* and *T*_1_ are given in Table 1 in the main text. Function [11] 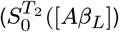 was smoothed in the same way.

**Figure S4.**
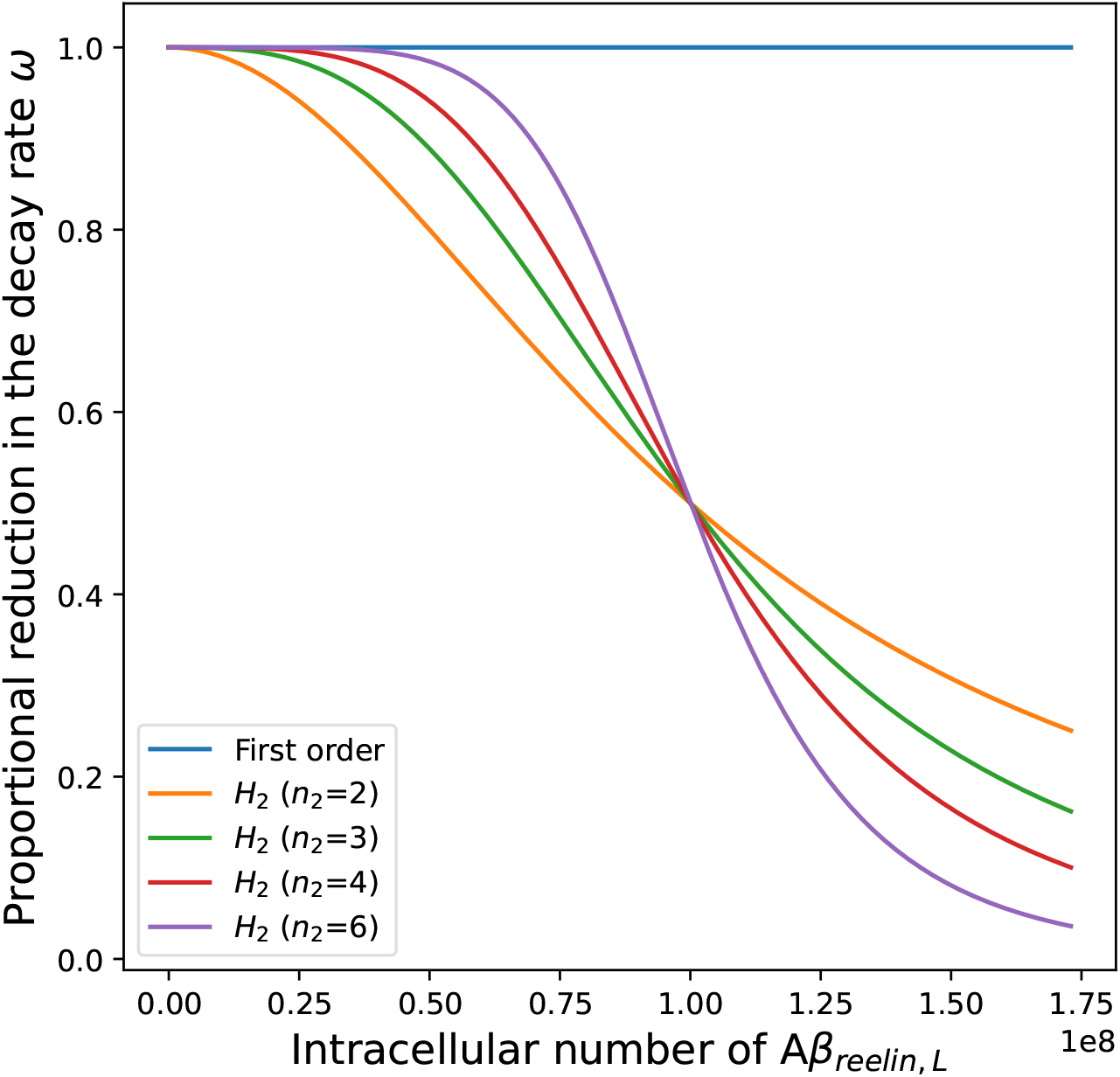
Examples of function [9] (*H*_2_([*Aβ*_*reelin,L*_], *θ*_2_, *n*_2_)) that is part of Eq. [7]. The blue line refers to a first order process, i.e. the degradation rate is given by *ω*[*Aβ*_*reelin,L*_]. *θ*_2_ = 1×10^8^. Since we assume the existence of a moderate sigmoidal doseresponse relationship, we chose to use the green inverse Hill function graph with *n*_2_ = 3.

**Figure S5.**
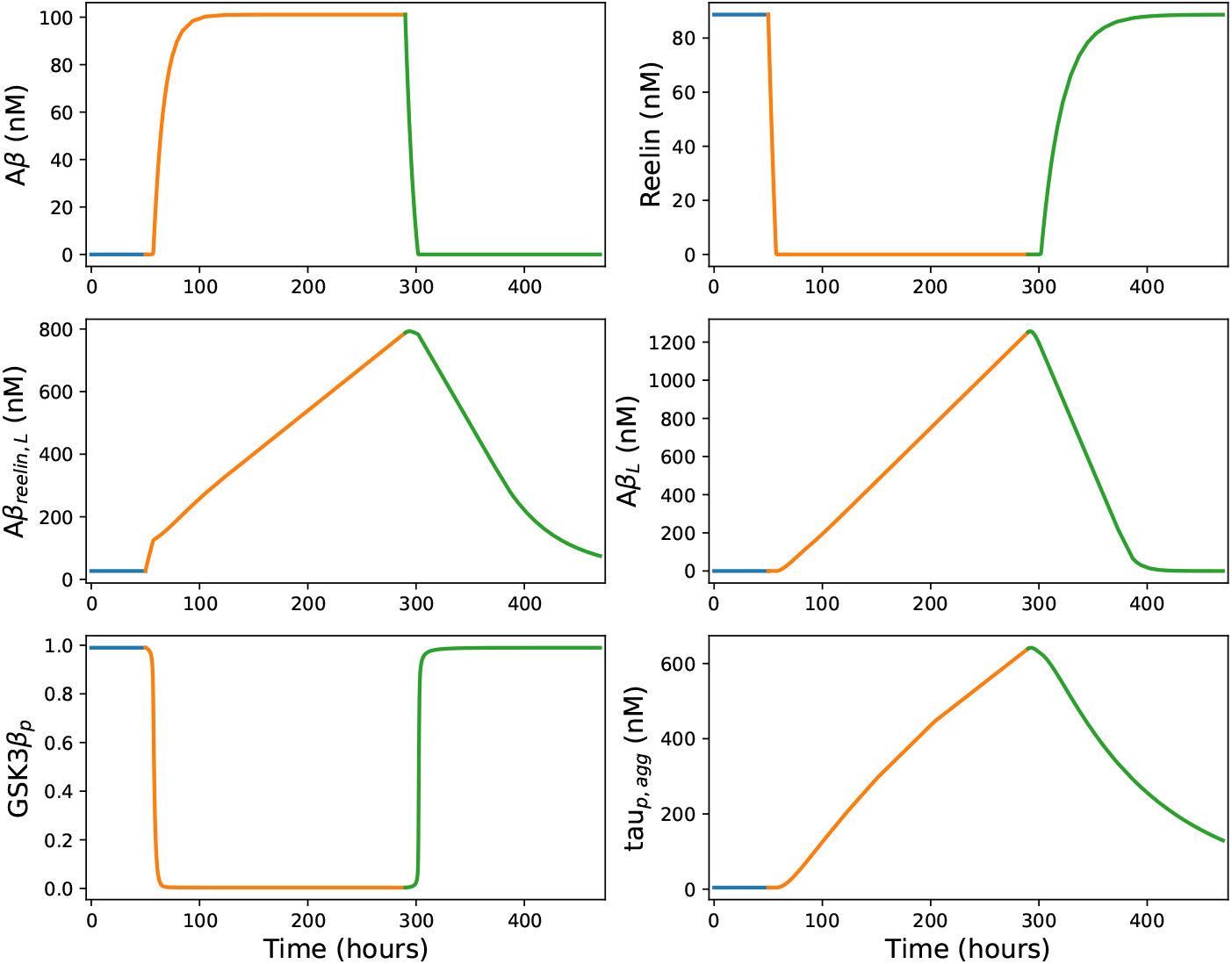
Short-term dynamic response to a temporary inflammation event in an LR neuron. The parameter values are given in Table 1 in the main text.

**Figure S6.**
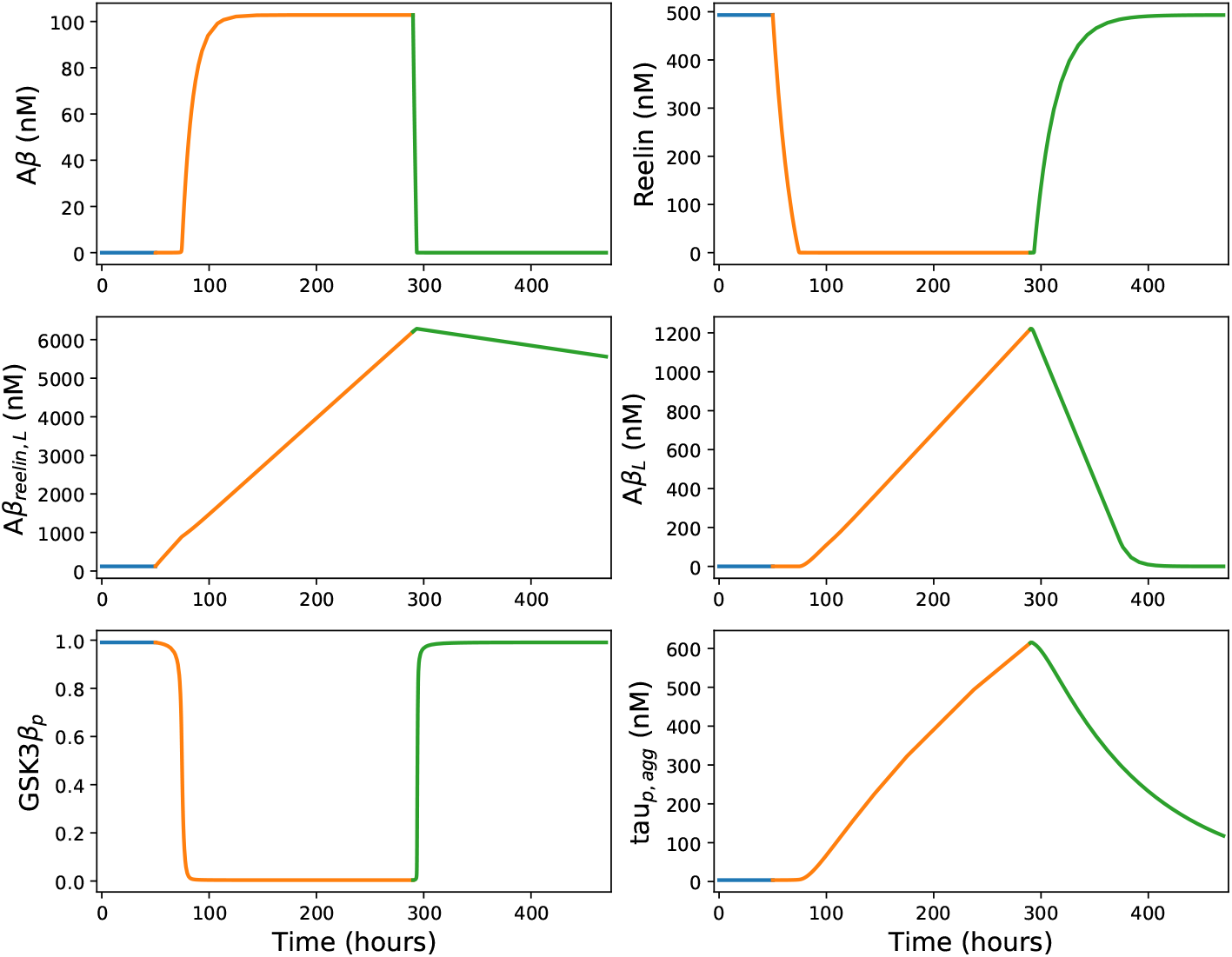
Short-term dynamic response to a temporary inflammation event in a Re^+^alECLII neuron. The parameter values are given in Table 1 in the main text.

**Figure S7.**
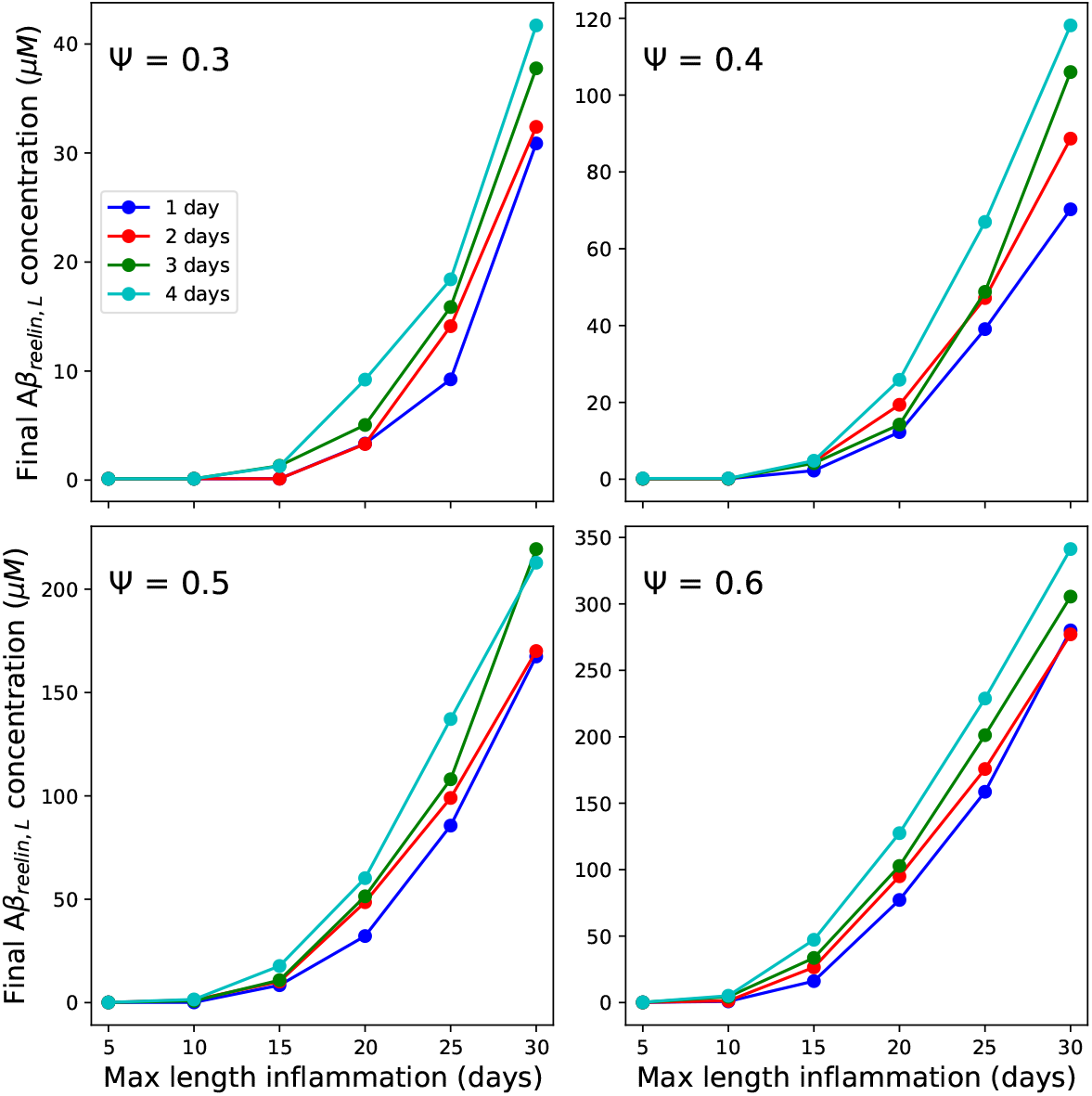
The impact of varying the parameters Ψ, *D*_*min*_ (see legend upper left panel) and *D*_*max*_ (see x-axis tick labels) underlying the inflammation event vector *I* on the endpoint concentration of *Aβ*_*reelin,L*_ in Re^+^alECII neurons.

**Figure S8.**
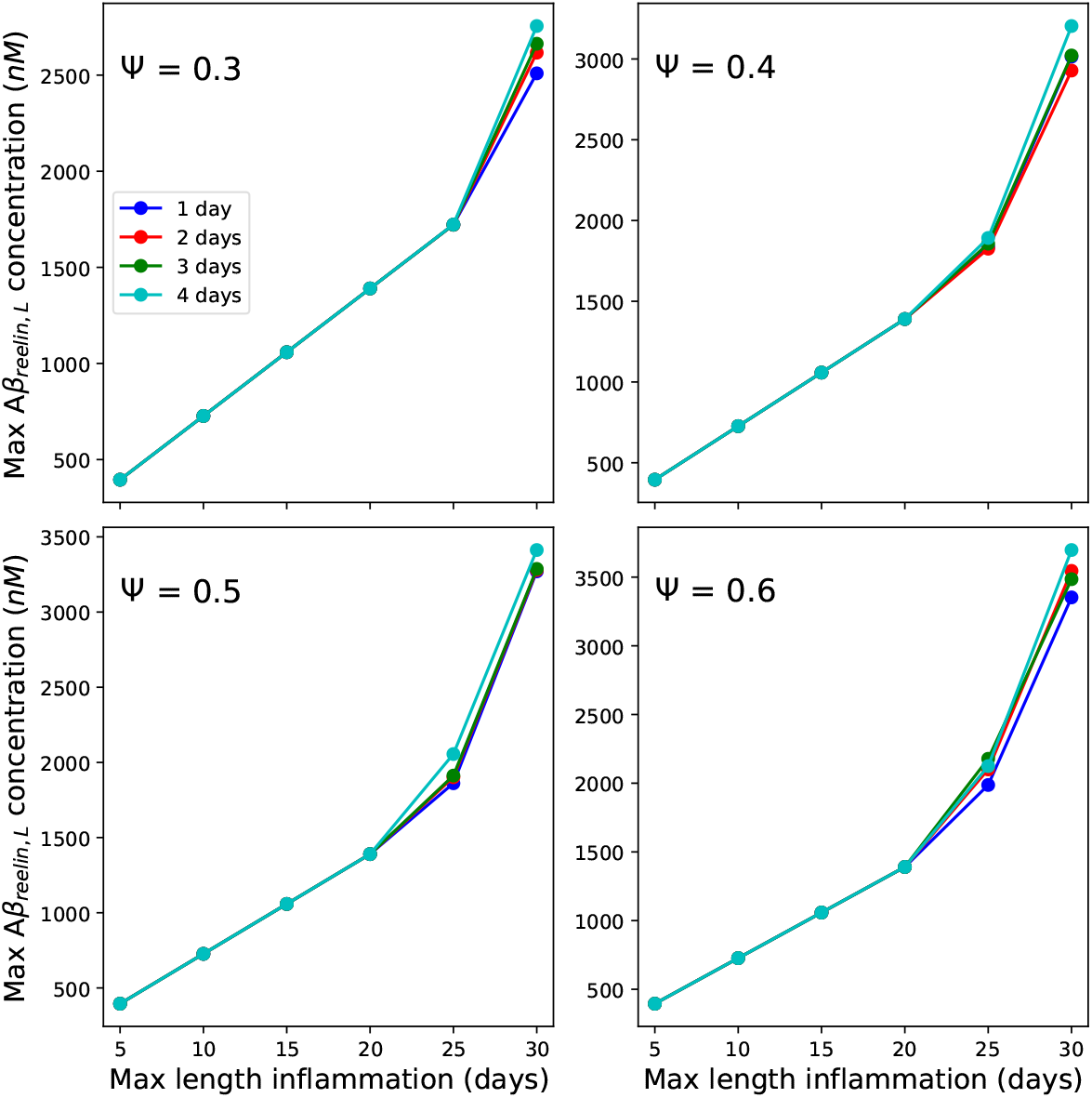
The impact of varying the parameters Ψ, *D*_*min*_ (see legend upper left panel) and *D*_*max*_ (see x-axis tick labels) underlying the inflammation event vector *I* on the maximum concentration of *Aβ*_*reelin,L*_ in LR neurons.

**Figure S9.**
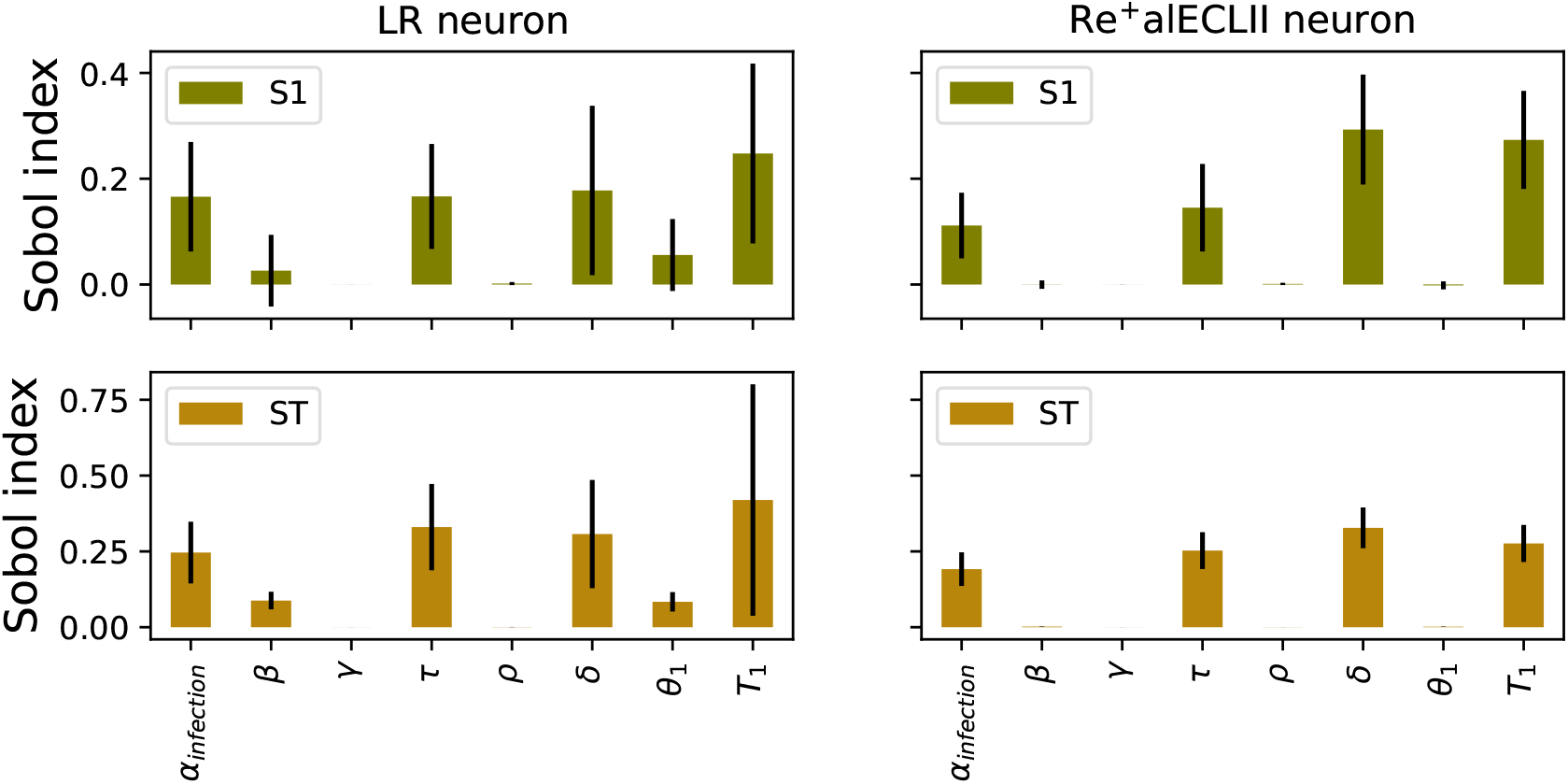
Sobol first order (S1) and total order (ST) indices with 95% confidence intervals for *Aβ*_*reelin,L*_ endpoint concentration-related parameters of LR and Re^+^alECII neurons (Saltelli et al., 2010), using the open source Python library SALib v 1.47 (Herman and Usher, 2017). The S1 index describes the influence of a specific parameter on the endpoint value of *Aβ*_*reelin,L*_ when all other parameter values remain at their nominal values. The ST index describes the influence of a given parameter and its interactions with other parameters. The analysis suggests that in LR neurons, the variation in *α*_*infection*_ has a much larger impact than *τ* and the degradation parameters *β, δ, θ*_1_ and *T*_1_ (Fig. S7, left panel column), while in Re^+^alECII neurons, the endpoint concentration of *Aβ*_*reelin,L*_ is more sensitive to the variation in *τ, δ*, and *T*_1_ (Fig. S9, right panel column). This implies that the predicted accumulation of *Aβ*_*reelin,L*_ can be directly linked to the observation that the reelin production rate is much higher in Re^+^alECII neurons than in LR neurons (Kobro-Flatmoen and Omholt, 2025), and that it depends very much on the degradation dynamics of *Aβ*_*reelin,L*_. That is, either the degradation rate must be very low in general (*δ*) or the degradation machinery becomes saturated above a given concentration (*T*_1_).

**Figure S10.**
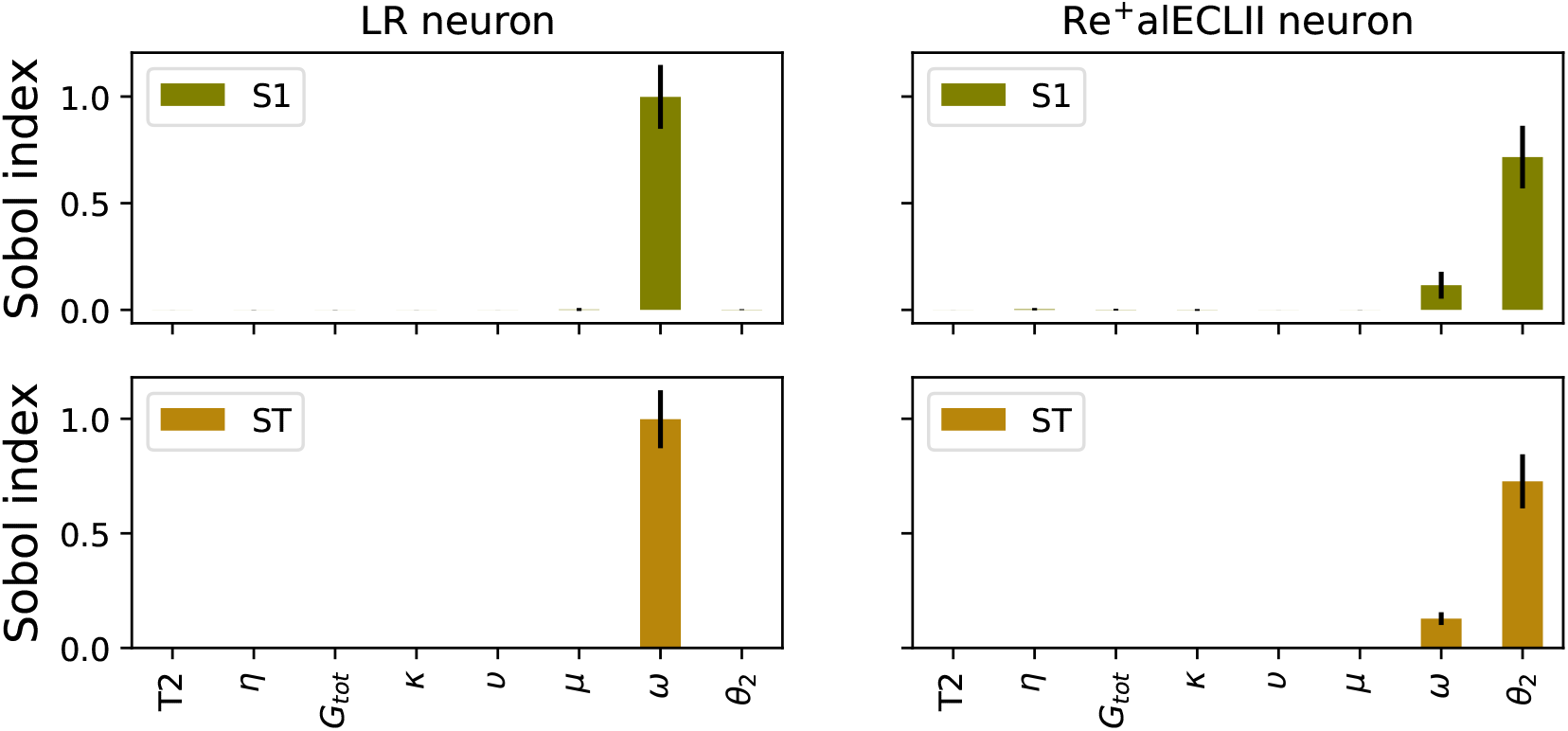
Sobol first order (S1) and total order (ST) indices with 95% confidence intervals for *tau*_*p,agg*_ endpoint concentration-related parameters of LR and Re^+^alECII neurons. The parameters of Eqs. [1]-[3] were fixed at their nominal values, Eq. [4] was ignored, and the parameter *ϕ* in Eq. [7] was fixed to 1.0. *n*_2_ was fixed to 3, while the remaining seven parameters in Eqs. [5]-[7] were allowed to vary between ± 50% of their nominal values. The number of generated parameter sets was 4608, with a discrepancy of 0.0007. The analysis followed the same procedure as described for Fig. S7. The Sobol analysis shows that the *tau*_*p,agg*_ endpoint concentration is predominantly sensitive to variation in the threshold value (*θ*2) where accumulated *Aβ*_*reelin,L*_ is assumed to begin to severely inhibit the degradation of aggregation-prone truncated forms of p-tau.

**Figure S11.**
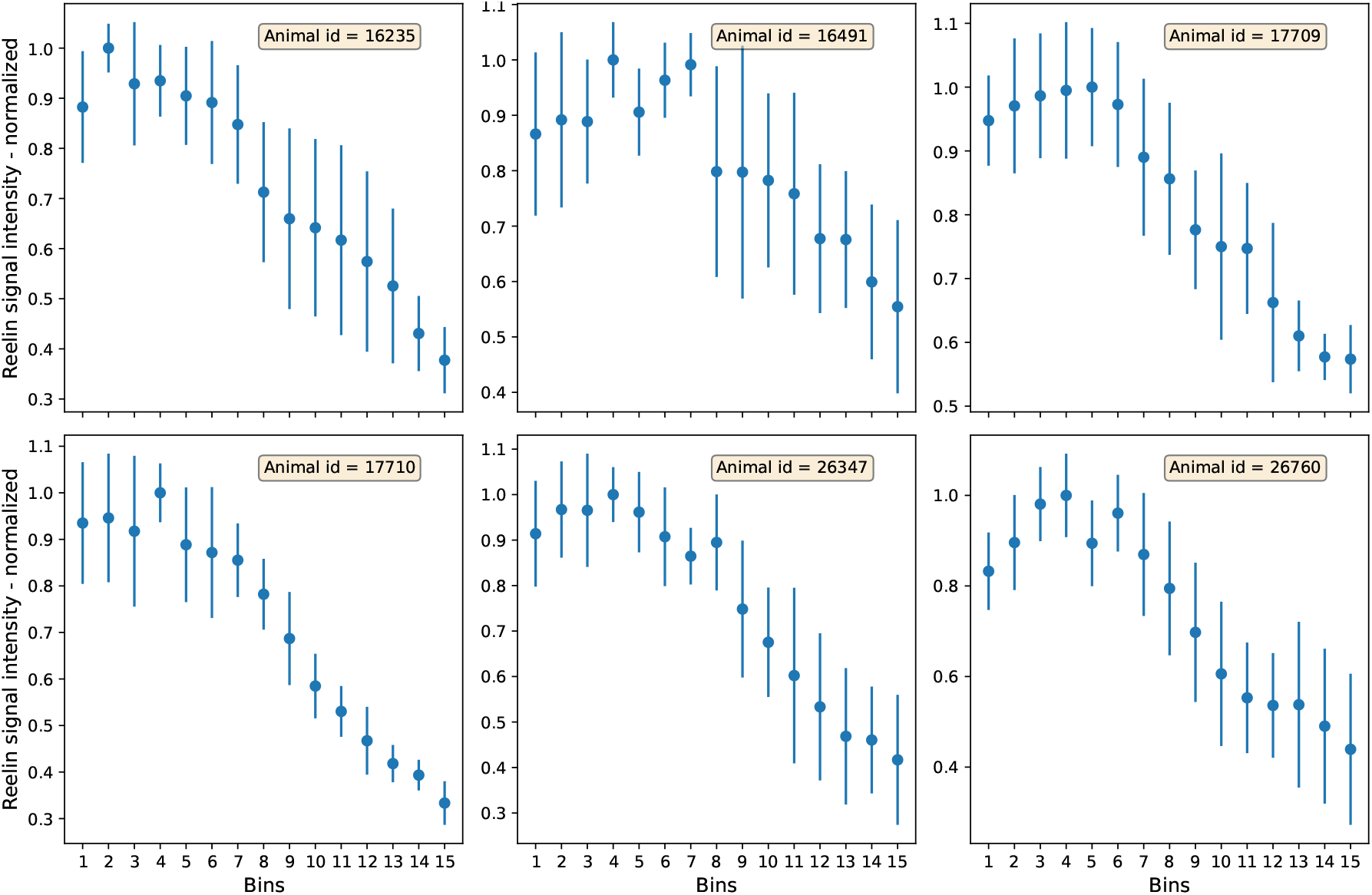
Reelin expression gradient in ECLII reelin expressing neurons in wild type Wistar rats. ECLII reelin signal intensity was measured from the lateral extreme (bordering the rhinal fissure) and successively more medial positions, up to and including the point at which the signal has the lowest intensity, which corresponds approximately to the border that separates the cytoarchitectonically defined lateral entorhinal cortex from the medial entorhinal cortex. The measurements were segregated into 15 equally sized bins. Methods in brief: Transcardial perfusions and tissue processing were done as described (Kobro-Flatmoen et al., 2023). Immunolabeling (40 um sections) included anti-reelin (1:1000; G10 clone; Merk, Cat# MAB5364; RRID: AB_2179313) with secondary antibody Alexa 635 Goat anti-mouse (1:500; ThermoFisher, Cat# A-31574). Densitometric measurements of reelin signal were done in QuPath (version 0.5.0), where we delineated ECLII and calibrated the ‘cell detection’ feature to detect reelin expressing neuronal somata. We used six animals (three 3-month-olds (IDs: 16235, 16491, 17709) and three 12-month-olds (IDs: 17710, 26347, 26760)). From each animal, to get optimal coverage of the lateral-to-medial extent, we used seven roughly equally spaced sections contained within Bregma -6.12 mm to -8.16 mm.

**Figure S12.**
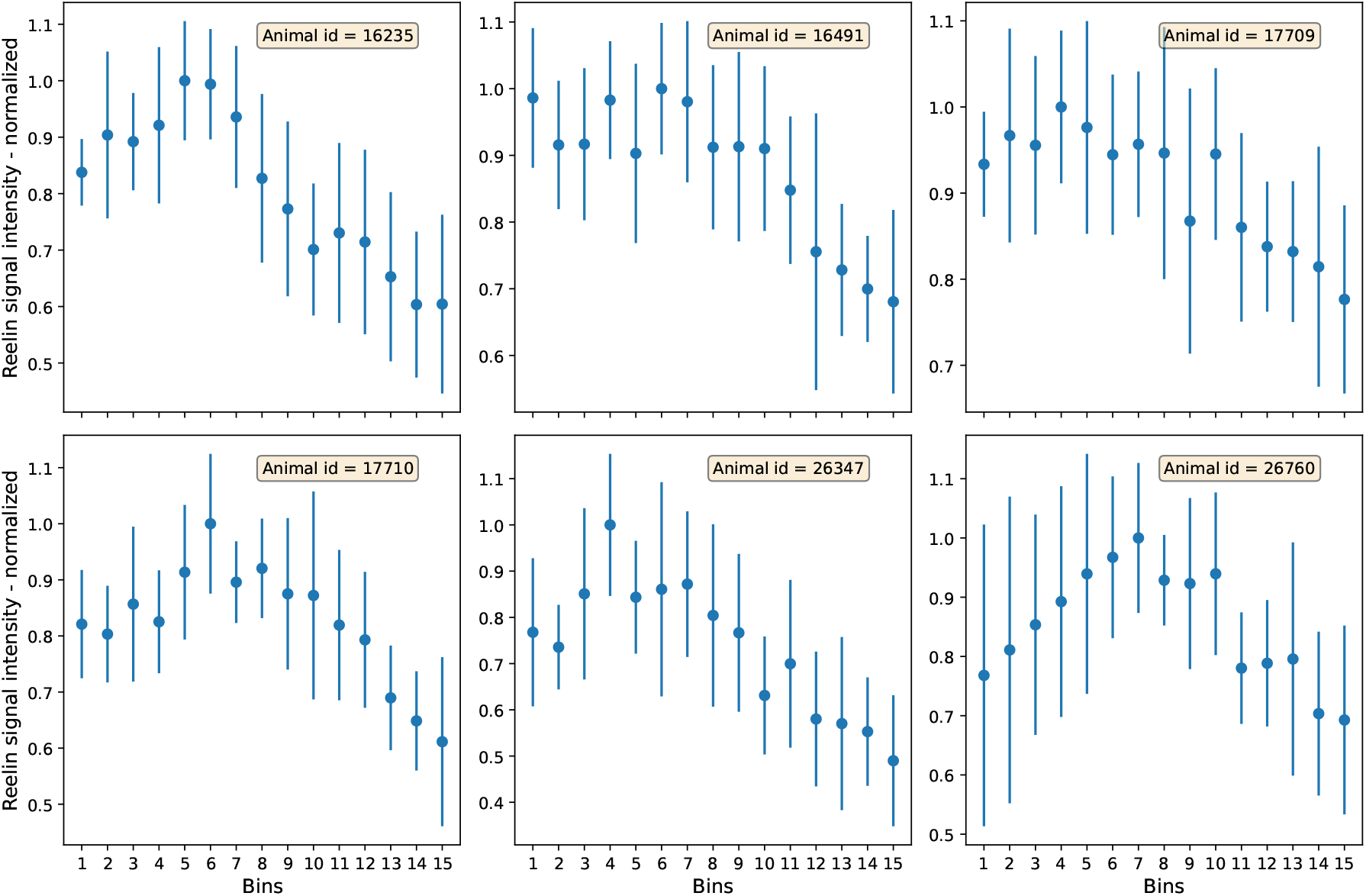
Lateromedial reelin signal intensity in ECLIII of 3 (IDs: 16235, 16491, 17709) or 12 (IDs: 17710, 26347, 26760) month old wild type rats. Each data point is the mean value of 7 tissue slices, normalized by the maximum signal value across the 15 bins. Experimental protocol identical with that of ECLII (see caption to Fig. S11).

**Figure S13.**
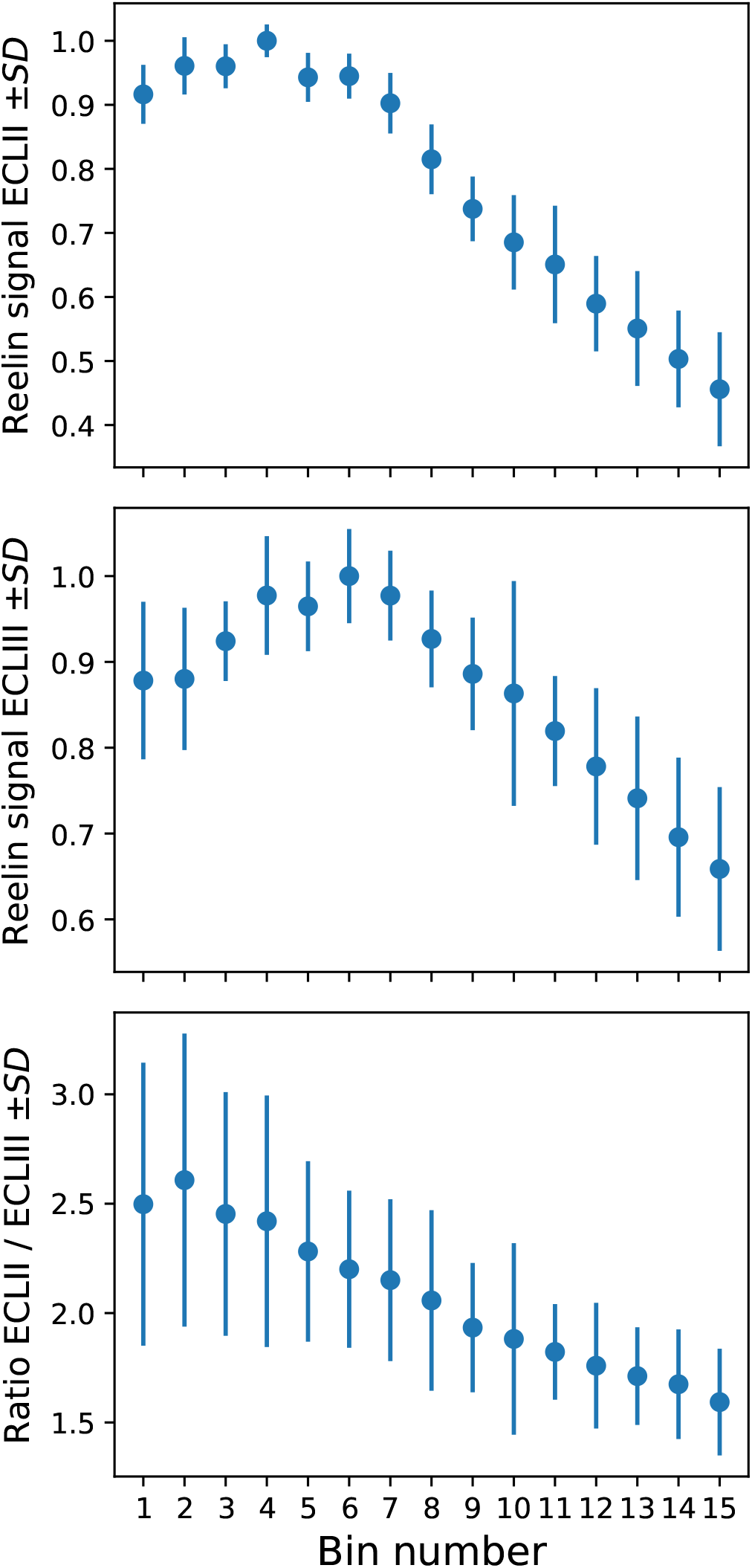
Upper and middle panels: Normalized lateromedial reelin signal intensity across the 15 bins in ECLII and ECLIII averaged over all seven tissue sections and all six animals. Lower panel: The lateromedial reelin signal intensity ratio of ECLII and ECLIII averaged over all seven tissue sections and all six animals.

